# Do Protein Language Models Learn Phylogeny?

**DOI:** 10.1101/2024.09.23.614642

**Authors:** Sanjana Tule, Gabriel Foley, Mikael Bodén

## Abstract

Deep machine learning demonstrates a capacity to uncover evolutionary relationships directly from protein sequences, in effect internalising notions inherent to classical phylogenetic tree inference. We connect these two paradigms by assessing the capacity of protein-based language models (pLMs) to discern phylogenetic relationships without being explicitly trained to do so. We evaluate ESM2, ProtTrans and MSA-Transformer relative to classical phylogenetic methods, while also considering sequence insertions and deletions (indels) across 114 Pfam datasets. The largest ESM2 model tends to outperform other pLMs (including the multimodal ESM3) by recovering phylogenetic relationships among homologous protein sequences in both low- and high-gap settings. pLMs agree with conventional phylogenetic methods in general, but more so for protein families with fewer implied indels, highlighting indels as a key factor differentiating classical phylogenetics from pLMs. We find that pLMs preferentially capture broader as opposed to finer evolutionary relationships within a specific protein family, where ESM2 has a sweet spot for highly divergent sequences, at remote distance. Less than 10% of neurons are sufficient to broadly recapitulate classical phylogenetic distances; when used in isolation the difference between the paradigms is further diminished. We show these neurons are polysemantic, shared among different homologous families but never fully overlapping. We highlight the potential of ESM2 as a complementary tool for phylogenetic analysis, especially when extending to remote homologs that are difficult to align and imply complex histories of insertions and deletions.

## 1 Introduction

For half a century, researchers have inferred phylogenetic “gene” trees from homologous protein sequences as a means to investigate protein function and evolution. This inference relies on an accurate multiple sequence alignment (MSA) of the proteins under study and involves statistical sequence modeling techniques such as Maximum-Likelihood and Bayesian statistics [1,2].Assumptions about evolutionary processes figure in the form of the Markovian substitution rate models such as Le-Gascuel (LG) [3]and the assumption of site independence. To mention but a few applications, this “classical” phylogenetic approach is used to improve the quality of an MSA [4, 5],to understand protein function, [6, 7] and for imagining ancient and novel proteins [8, 9].

Protein language models (pLMs) analyse vast amounts of proteins to uncover (non-linear) “language” patterns embedded in amino acid sequences, that in turn can predict structure and function. pLMs have been shown to detect remote homology [10, 11], discover evolutionary relationships within protein families [12, 13], align sequences in the “twilight zone” [14, 15], predict variant function [16], and guide directed evolution experiments [17, 18].

Phylogenetic analysis (with a universal substitution rate model) usually centres on a single family with a single origin, while pLMs are truly universal in the sense they project a protein sequence (or alignment of sequences in the case of the MSA-Transformer) from *any* family to a multi-dimensional, dense numerical vector space of sequence embeddings. We set out to examine the extent to which this space captures phylogenetic signals for all homologous protein sequences to understand the overlap and differences of the two paradigms. Such examination can in turn determine if these paradigms are two sides of the same coin, or if they complement one another by resolving questions that the other approach struggles with.

Lupo et al. [19]showed that MSA-Transformer [20]allocates without explicit guidance an internal “attention head” that recapitulates the Hamming distance between aligned protein sequences. Here, we examine the extent to what evolutionary relationships amongst protein sequences (as recovered by conventional phylogenetics) are encoded by pLMs trained on large protein sequence corpora and available via querying that pLM for any *pair* of protein sequences. We ask:

- How well does a sequence embedding of pre-trained pLMs recover the evolutionary relationship inferred by classical evolutionary models? What (if any) difference can be discerned between the embedding of an MSA-based pLM and that of a single-sequence based pLMs? Does the implied presence of insertions and deletions (indels) influence either assessment?
- How similar are the local homologs in pLM embedding spaces to the classical evolutionary models?
- Is the evolutionary relationship recovered by pLMs subject to distance?
- When demonstrated for pairs of homologous proteins, how is their evolutionary relationship encoded and organised in the embedding space? Is it localised or distributed?

We seek answers to the above questions for models trained on (1) unaligned and (2) aligned protein sequences, from 114 Pfam datasets. For unaligned proteins, we utilise single-sequence models ESM1, ESM2 [21], and ProtTrans [22]; for aligned proteins, we utilise the MSA-Transformer [20].Additionally, we address if single-sequence models trained on sequences alone (unimodal) differ significantly (in terms of the above questions) to models trained on multimodal input (e.g., structure, function), such as ESM3 [23] and ProstT5 [24].

We draw the following conclusions. With few exceptions, unaligned, single-sequence models recover phylogenetic relationships more accurately than MSA-based models, for both low- and high-gap sequence inputs, with ESM2 showing the best performance overall. Unsurprisingly, we find that pLMs perform more reliably on protein families with fewer implied indels compared to those with high indels. MSA-Transformer based pLM representations show vastly different performance in low-gap and high-gap settings, with performance in low-gap settings comparable to that of single-sequence models. When evaluated for sensitivity at different evolutionary distances, relationships uncovered by ESM2 agree more closely with classical phylogenetic analysis when sequences are divergent, at greater distances. Single-sequence pLMs tend to use layers quite differently from multiple-sequence pLMs, with early layers in the former class learning basic properties of sequence components before their relationships. MSA-Transformers appear to encode higher-level organisation of relationships early, implied by an attention-based architecture deconvoluting the input MSA. To understand different models’ capacity, we probe the output neurons and find that a small subset (*<* 10%) is responsible for encoding evolutionary relationships. Interestingly, these neurons overlap across different protein families but are never fully shared, highlighting their polysemantic nature.

## 2 Preliminaries

### 2.1 Experimental Setup

#### 2.1.1 Datasets

In this study, we use 114 MSA-and-tree datasets from Pfam [25]stratified into low-gap and high-gap MSAs to investigate how and how well different pLMs recapitulate implied evolutionary distances. Pfam provides a comprehensive collection of protein families and domains, which are curated and classified according to their evolutionary relationships, structure, and function.

We extract datasets from Pfam based on the gap percentage of the MSA to analyse the impact of indels on evolutionary relationship captured by different pLMs. The low-gap dataset from Pfam comprises of 30 protein families and 30 protein domains, each exhibiting a gap percentage of 20% or less, with a minimum of 200 total sequences and lengths greater than or equal to 70. The high-gap dataset consists of 27 protein families and 27 protein domains with a gap percentage of 70% or greater, each with with 200 or more sequences of lengths greater than or equal to 70. Gap percentage is calculated by dividing the total number of gap positions by the total number of positions in the MSA. The distributions of gaps, numbers, and lengths of sequences found in MSAs are shown in Supplementary Figure S1. Full list of datasets are in Supplementary Table S1.

#### 2.1.2 pLM representations

We initially probe five different unimodal pLMs (see Table 1) to inform a selection for further analysis. For each pLM we retrieve embeddings from different layers. The single-sequence models esm2 t48 15B UR50D, esm1b t33 650M UR50S, prot t5 xxl uniref50, and prot t5 xxl BFD process protein sequences (with gaps removed) in batches, while the esm msa1b t12 100M UR50S (MSA-Transformer) model takes a whole MSA in one go. Embeddings from pLMs can capture rich semantic information for each protein sequence; here, we use the mean embedding of a layer as the pLM representation for a protein sequence.

**Table 1.**
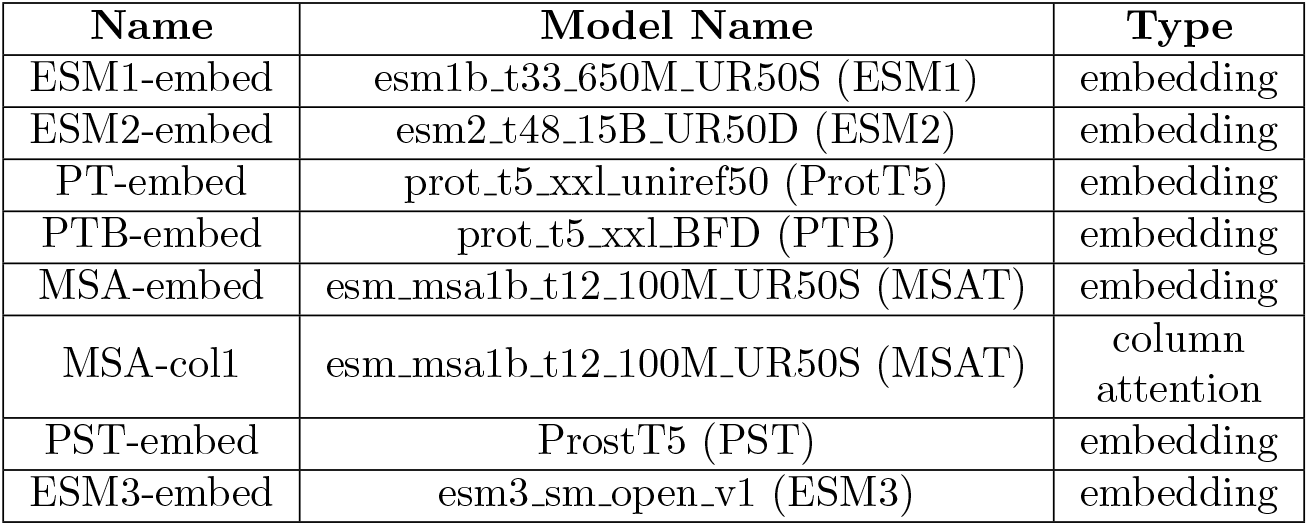
Different pLM representations

| Name | Model Name | Type |
| --- | --- | --- |
| ESM1-embed | esm1b_t33_650M_UR50S (ESM1) | embedding |
| ESM2-embed | esm2_t48_15B_UR50D (ESM2) | embedding |
| PT-embed | prot_t5_xxl_uniref50 (ProtT5) | embedding |
| PTB-embed | prot_t5_xxl_BFD (PTB) | embedding |
| MSA-embed | esm_msa1b_t12_100M_UR50S (MSAT) | embedding |
| MSA-col1 | esm_msa1b_t12_100M_UR50S (MSAT) | column<br>attention |
| PST-embed | ProstT5 (PST) | embedding |
| ESM3-embed | esm3_sm_open_v1 (ESM3) | embedding |

Formally, for a given protein sequence *s* defined by its sequence of *l*_*s*_ amino acids, we obtain corresponding latent representation 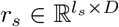 from each single-sequence pLM model. Each amino acid of protein sequence is mapped into ℝ_*D*_. From latent representation 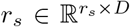, we obtain protein-level representation *e*_*s*_ ∈ ℝ_1*×D*_ by applying average pooling layer on the amino-acid-level features over sequence length given b

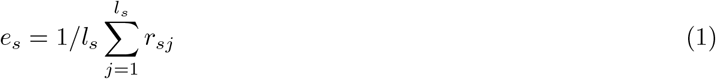

Following Lupo and colleagues [19],we use a comparable average pooling technique for the MSA-Transformer to extract sequence embeddings, but with a fixed length for each protein sequence in the MSA. Additionally, we extract the column attention heads. The various pLM representations are outlined in Table 1 along with the name we use in this paper.

### 2.2 Methods

#### 2.2.1 Representational Similarity Analysis (RSA)

RSA originates from cognitive neuroscience and is used to investigate the similarity of heterogeneous representational spaces for different stimuli or conditions [26, 27].The different stimuli under consideration are described in the next section.

In our application, we pass the individual Pfam datasets through each pLM and extract their latent representation. We adopt the Euclidean distance as the kernel for calculating the pairwise similarity for all sequences when presented to the pLM. We refer to this similarity matrix as the pLM matrix (**M**_pLM_) and in many cases it will be specific to the “layer” of the neural network.

For the column attention of MSA-Transformer, we average the attention values across the sequence length. To ensure symmetry, we transpose the matrix and add the attention values between sequences. While the operation on column attention could be reversed by summing up the attention per site of the sequence and then averaging it, we follow the method outlined in [19].

We use the (additive) branch distance in phylogenetic tree to quantify similarity between any pair of protein sequences as per classical phylogenetics. We use FastTree [2]with the LG evolutionary model [3]to construct phylogenetic trees for each Pfam dataset. We compute all pairwise branch distances using the ete3 Python package [28]as the average across ten trees, each inferred for a dataset using different random seeds (see Supplementary Figure S2). We refer to the average distance matrix as the LG matrix (**M**_LG_) ; this matrix is used to gauge if matrices generated from pLMs reflect a signal that is captured by an evolutionary model, which in turn depends on an accurate sequence alignment.

Finally, we consider a baseline amino acid composition space. Using the so-called onehot encoding, every amino acid in a protein sequence is presented as a binary, unit-length vector. Averaging the onehot vector across all positions yields a distribution of amino acids, representing the sequence insensitive to the order of amino acids. We use the Euclidean distance as the kernel to represent pairwise similarity between onehot sequence representations; we refer this as the onehot matrix (**M**_OH_) and use it to compare the extent to which other matrices capture basic compositional information, which in turn does *not* depend on accurate alignment.

To compare representational spaces, we define an Evolutionary Similarity Score (*ESS*). For a Pfam dataset *d*, we compute the similarity matrices **M**_pLM_ and **M**_LG_, and calculate the similarity *S* using Spearman’s rank correlation coefficient (*ρ*) and Pearson’s correlation coefficient (*r*) as follows:

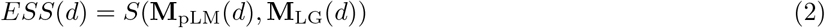

Thus we refer to *ESS* calculated using *ρ* and *r* as *ESS*_*ρ*_ and *ESS*_*r*_ respectively. In our application, we consider four different stimuli for RSA.

- Stimuli 1: We evaluate the internal layers of pLMs for amino acid composition, via the onehot matrix, for each Pfam dataset, to quantify the sensitivity to conserved content without consideration to position.
- Stimuli 2: We consider the evolutionary distances across the phylogenetic tree for each Pfam dataset, to assess the ability of pLMs to capture pairwise evolutionary relationships.
- Stimuli 3: We consider ten evenly spaced sorted protein sequences to a reference sequence within the Pfam dataset, to establish if pLMs capture evolutionary relationships at broad evolutionary timescale.
- Stimuli 4: We consider the ten closest sequences to a reference sequence within the Pfam dataset, to assess the resolution of pLMs at a finer timescale.
- Stimuli 5: We consider systematically shuffled sequences (aligned and non-aligned), to establish the standard of performance that is explained by chance. For each protein sequence, two sites are randomly selected and their amino acids are swapped. This procedure is applied for 80% sites of the total sequence length, for each protein sequence. In analyses involving the ESM2 and ProtTrans models, shuffling is conducted on non-aligned protein sequences. For the MSA-Transformer, shuffling is conducted on aligned sequences, including gap characters.

#### 2.2.2 Local homolog similarity (LHS)

We compare the local neighborhoods in the phylogenetic tree and the pLM embedding space with respect to a reference sequence (*r*_*s*_). For each *r*_*s*_, we identify the k-nearest sequences (*K*_*LG*_) based on evolutionary distance in the phylogenetic tree, forming a sequence set. Similarly, for the same reference sequence, we compute the k-nearest sequences (*K*_*pLM*_) in each pLM embedding space using Euclidean distance. The similarity (*J*) between these two sets, *K*_*LG*_ and *K*_*pLM*_, is measured using the Jaccard similarity coefficient, defined as the intersection over union of the two sets, as shown below:

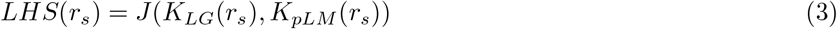

#### 2.2.3 Explainable-AI methods probe how pLMs encode homology

Probing classifiers are used in the field of Explainable-AI to identify and analyse individual or sets of neurons, to in turn understand what information the network has learned and how it is encoded.

First, we use pLM representations to carry out an “evolution” probe using an Elastic-Net regression model. We use a linear model because the learned weights can be used to measure the importance of predictor variables. Elastic-Net regression offers Lasso (*L*1) and Ridge (*L*2) regularisation; it is most suitable in our case as neurons are multivariate in nature and work in groups. The model is trained to predict the evolutionary distance between two protein sequences based on the absolute difference between their pLM representations.

Formally, lets consider a pLM *P* and for a Pfam dataset *F* = *{s*_1_, …, *s*_*N*_ *}* with *N* protein sequences and corresponding LG matrix *L* of size *N ×N*. We map each protein sequence *s*_*i*_ in *F* to the output latent representations: 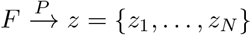. For every pair of *s, s* with *i, j* ∈ *N* and *i* ≠ *j*, we calculate |*z*_*i*_ − *z*_*j*_ | and train the regression probe to predict *L*[*s*_*i*_][*s*_*j*_]. The model is trained to minimise the following loss function,

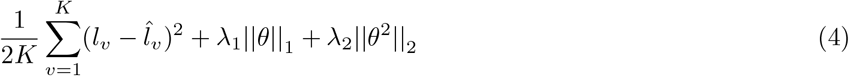

where *K* is the number of training samples and *l* is the evolutionary distance as per the LG matrix and *Î* is the evolutionary distance predicted by the regressor. The weights *θ* ∈ *ℝ*_*D*_ are learned by the regressor and *D* is the dimensionality of the latent representations *z*_*i*_. The terms *λ*_1_ and *λ*_2_ are the *L*1 and *L*2 regularisation parameters, which are optimised for each dataset using cross validation.

Given the trained regressor, we consider the weights assigned to each neuron in |*z*_*i*_ − z_j_| as a measure of their importance with respect to the evolutionary information between protein sequences. We arrange the absolute value of the neurons in the descending order and consider the top neurons constituting 25% of the weights of all neurons for analysis. Similarly, we also consider the bottom neurons constituting 25% of the weight of all neurons.

## 3 Results

We processed each Pfam dataset through each pLM to obtain their representation (see Table 1),including all 30 low-gap protein domains and 30 families. Some high-gap datasets failed for the MSA-Transformer due to its maximum input sequence length and depth of 1024, in turn limiting the analysis to 27 high-gap protein domains and 21 high-gap protein families. (Single-sequence pLMs also have a length limit but accept unaligned sequences, in effect permitting longer sequences.)

### 3.1 Single-sequence pLMs learn the amino acid composition of proteins in the initial layers

We first asked what internal layers of pLMs contain compositional information of protein sequences. We used RSA between the **M**_OH_, which compares the amino acid distributions between pairs of sequences, and each of the pLM matrices (**M**_pLM_), thereby assessing the *ESS*_*ρ*_ (Spearman’s rank) and *ESS*_*r*_ (Pearson’s correlation coefficient) across all internal layers of the pLMs. Figure 1(a), 1(b), 1(c) shows the *ESS*_*ρ*_ but similar trends were observed for *ESS*_*r*_ (see Supplementary Figure S3).

**Figure 1.**
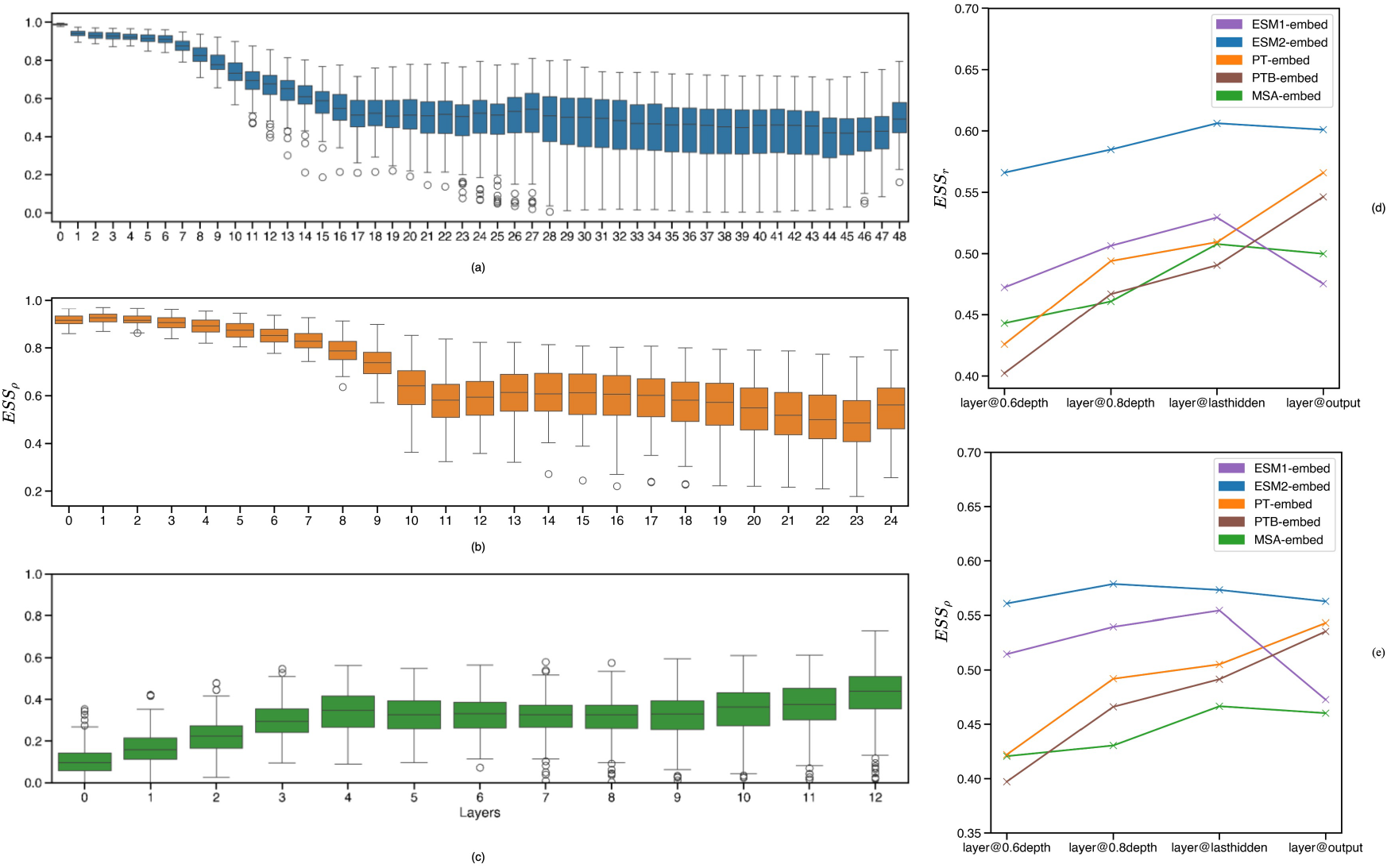
Evolutionary similarity score (*ESS*_*ρ*_; y-axis) between layer-specific pLM matrix (x-axis) and onehot matrix. Boxplots are based on 114 Pfam datasets for ESM2-embed (a) and PT-embed (b). In case of MSA-Transformer (c), boxplots are based on 100 Pfam entries. Line plot (d) compares *ESS*_*r*_ (y-axis) and line plot (E) compares *ESS*_*ρ*_ (y-axis) for pLMs based on matrices at different layer depths (x-axis).

The *ESS*_*ρ*_ peaks in the initial layers of the network for all single-sequence pLMs, then gradually decreases in the middle before plateauing in later layers. This observation is consistent with a top-down learning process that encodes basic but general organisation (of amino acids) in early layers, to then reflect specific conservation present in smaller groups of proteins in later layers. We hypothesise that learning incorporates higher-order dependencies in later layers amongst the basic components encoded in those preceding; we test this in the next section.

Interestingly, the MSA-Transformer exhibits a trend distinct to that of single-sequence models, potentially aided by a pre-aligned input. It demonstrates a gradual increase in *ESS*_*ρ*_ in the later layers; this increase is consistent with a bottom-up encoding: early layers reflect relationships that are specific to subsets of data, with later layers establishing how those relationships can be generalised across families. While layers render evolutionary information differently, all pLMs exhibit a degree of correlation with the **M**_OH_ in their later layers.

### 3.2 ESM2 agrees with evolutionary models for both gappy and non-gappy homologous protein sequences

To explore if pLMs reproduce relationships that are recovered using classical phylogenetic analysis, we use RSA between the LG matrix (**M**_LG_) and pLM matrices (**M**_pLM_) for a select number of layers within each pLM model.

Valerian and colleagues [29] suggest that pLM layers with a relative depth of 0.6 to 0.8 in relation to the total number of layers are optimal for identifying remote homology, while Rives and colleagues [10] submit that the last hidden layer does this best. With those studies and the previous analysis in mind, we extract embeddings at 0.6 depth, at 0.8 depth, the last “hidden” layer, and the output layer.

Across all pLM embeddings, later layers had the greatest correlation with the **M**_LG_, which here represents a classical phylogenetic distance matrix (Figure 1(d) and 1(e)).Relative to ESM1-embed, ESM2-embed has a consistently greater correlation with LG, with the last hidden layer or the output layer showing this most convincingly. MSA-embed had a comparatively lower correlation with LG, with stronger correlation in its last hidden, marginally higher than its output layer. Within the ProtTrans model family, PT-embed has a consistently greater correlation with the LG matrix than PTB-embed, again across all layers, with the output layer being the strongest for both. Based on these findings, we opt to proceed with ESM2, PT-embed and MSA-embed with the embeddings from their output layer for detailed analysis.

To put this into perspective of previous work, we include a column attention in layer one head five from the MSA-Transformer; henceforth, referred to as MSA-col1. Lupo et al. [19] demonstrated that this particular column attention recapitulates the Hamming distance between protein sequences. The final pLM representations are named and summarised in Table 2.

**Table 2.**
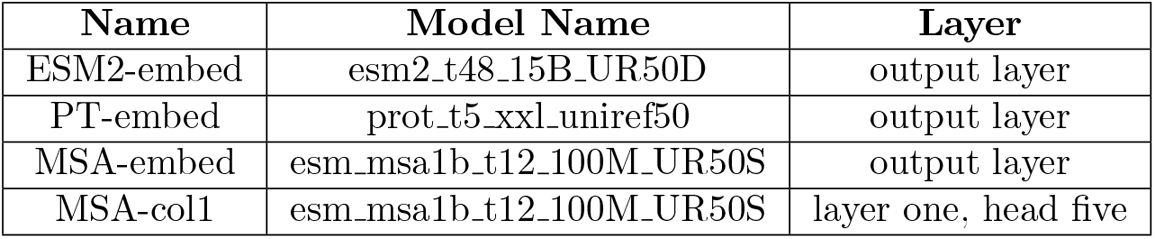
pLM representations for evolutionary probe

We investigate whether embeddings for homologous domains or entire protein sequences (as presented in families) are more (or less) suited to detect phylogenetic relationships. A two-sample Kolmogorov-Smirnov statistical test [30]for goodness of fit, comparing the *ESS* distribution between types of embeddings within the low-gap and high-gap categories, show no statistically significant differences in *ESS* (both *ESS*_*ρ*_ and *ESS*_*r*_). This, in turn, justified not separating their analyses below.

There is some variability in *ESS* between datasets for different pLM matrices (Table 2) against the LG matrix (Figure 2(a) and 2(b)). Notable is the low *ESS* for MSA-Transformer representations in the high-gap datasets. Given that the MSA-Transformer requires the input to be an MSA (like classical phylogenetic analyses, and by extension LG matrix), we anticipated it to outperform single-sequence pLMs, which do not require this information. Single-sequence pLMs have greater *ESS* overall than MSA-Transformer for high-gap datasets, with ESM2-embed topping both categories in terms of averages and a one-sided Wilcoxon signed-rank test (*P <* 0.05; both order and magnitude), except for *ESS*_*ρ*_ in the low-gap dataset when no significant difference is observed between ESM2-embed and MSA-col1. PT-embed ranks highest in the one-sided Wilcoxon signed-rank test (*P <* 0.05; both order and magnitude) against MSA-embed, except for *ESS*_*r*_ in high-gap datasets where no significant difference is observed. Similarly, PT-embed tops the test against MSA-col1, except for *ESS*_*ρ*_ in low-gap datasets where no significant difference is found. With MSA-embed and MSA-col1, the results are mixed.

**Figure 2.**
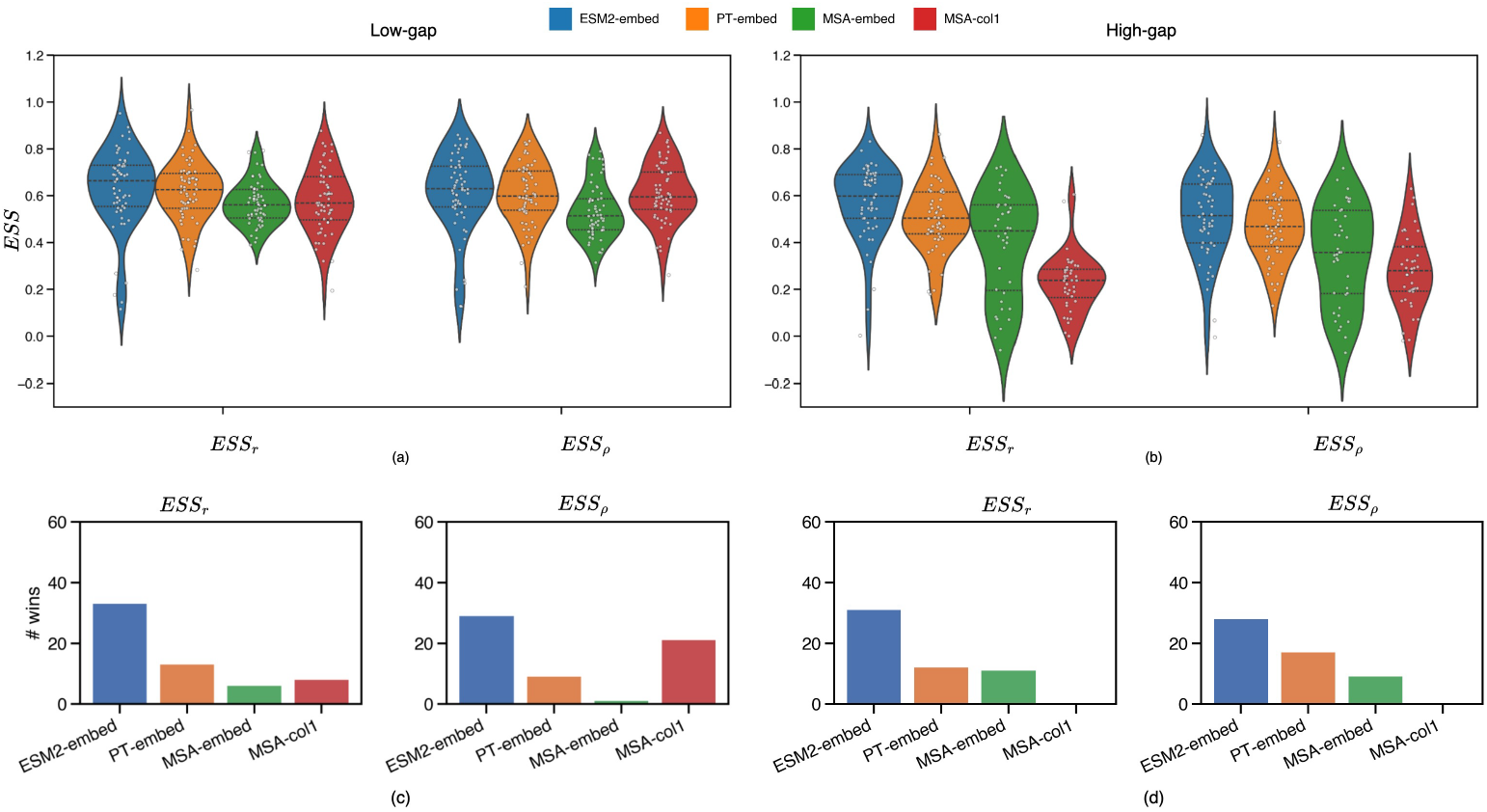
Violin plots for *ESS*_*ρ*_ and *ESS*_*r*_ for different pLM matrices against LG, across all applicable low-gap (a) and high-gap (b) datasets. Bar plots showing the number of datasets where the type of pLM has the greatest *ESS* across all applicable low-gap (c) and high-gap (d) datasets.

To ascertain how sensitive evaluations are to protein sequence integrity, we check the *ESS* signal against a randomly shuffled input sequences in each dataset. When shuffling 80% of sites, the *ESS* (to LG) for both ESM2-embed and PT-embed decreased significantly. Interestingly, the *ESS* of MSA-embed and MSA-col1 did not drop substantially. This phenomenon may be attributed to the model architecture, which incorporates row and column attention in each of its transformer blocks. The tied column attention could potentially still exert influence from the unshuffled column to the shuffled sequence in other sequences (Supplementary Figure S4). The one-sided Wilcoxon signed-rank test confirms that the *ESS* distribution of the normal dataset is significantly greater than the “80-shuffled” distribution for each pLM, but that pLMs vary in their sensitivity to the order of the amino acids.

When pLMs compete for each dataset (Figure 2(c) and 2(d)),ESM2-embed has the top *ESS*_*ρ*_ and *ESS*_*r*_ most times, in both gap categories. This superiority may partly be attributed to the sheer size of ESM2 with 48 transformer blocks and 5120 embedding dimension size. PT-embed ranks second overall, except in the *ESS*_*ρ*_ category for low-gap datasets, where MSA-col1 leads. Between MSA-embed and MSA-col1, MSA-col1 prevails in low-gap datasets for both *ESS* measures, while MSA-embed excels in high-gap datasets. This suggests that in high-gap datasets, the later layers of the MSA-Transformer play a crucial role in learning the evolutionary relationships between protein sequences, as the presence of high gaps muddies detection of homology.

There are select low-gap datasets (PF01948, PF13667, PF06421, PF08439, and PF00986) where ESM2-embed performs poorly (see narrow-tailed violin plots in Figure 2(a) for both *ESS*) while the MSA-Transformer exhibits higher *ESS* (*ESS*_*ρ*_ and *ESS*_*r*_) (≥ 0.5), in contrast to that of ESM2-embed (≤ 0.30). There are no distinct characteristics specific to these datasets, but the availability of MSAs in the low-gap datasets might offer additional context that aids in revealing evolutionary relationships between sequences, which may not be as critical for other datasets. Analysing the number of datasets where the LG correlation is worse (≤ 0.30 *ESS*), ESM2-embed has the highest number of “wins” at 5 in low-gap category, compared to PT-embed (1), MSA-Col1 (1) and MSA-Embed (0) for both *ESS* (Supplementary Table S2). For high-gap, MSA based pLM representations have highest “wins” (Figure 2(c) and 2(d)).

In general terms, the *ESS* is greater for low-gap compared to high-gap datasets across all pLM representations (*P <* 0.05; one-sided Wilcoxon rank-sum test), suggesting that gaps impede pLMs’ ability to recapitulate conventional phylogenetic analysis; it is noteworthy that classical phylogenetic analysis tends to ignore gaps, which could account for the differences observed in high-gap datasets.

### 3.3 MSA-Transformer uses “attention” to distinguish between evolutionarily similar and distinct proteins

Lupo et al. [19] probed column attention heads in MSA-Transformer, identifying one head that specifically correlates with the Hamming distance between sequences. We probed all heads across all Pfam datasets using the same approach, adapted for the LG matrix (see Section 2.2.1).As expected, layer one, head five, is *positively* correlated with evolutionary distance. Interestingly, the column attention heads in later layers of MSA-Transformer model typically have *negative ESS* (*ESS*_*ρ*_ and *ESS*_*r*_) for both low-gap and high-gap datasets, strongly exemplified by layer three, head 12 (henceforth MSA-col2). This flip in *ESS* implies that pairs of proteins which are “high” in attention, are similar in evolutionary distance, and (conversely) proteins which are “low” in attention, are evolutionary distinct. Figure 3 shows the distribution of *ESS*_*r*_ for MSA-col1 and MSA-col2 across all low-gap and high-gap datasets. Similar trends are observed for *ESS*_*ρ*_ (see Supplementary Figure S5).

**Figure 3.**
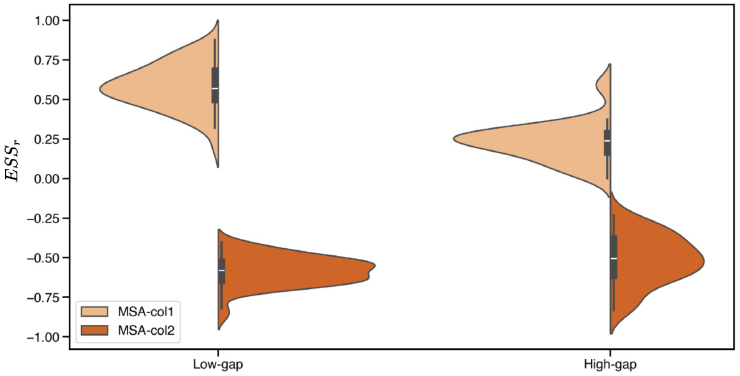
Distributions of *ESS*_*r*_ of select column attention heads in MSA-Transformer, across all applicable low-gap and high-gap datasets; the scores for MSA-col2 (layer three, head 12; dark orange) negatively mirrors MSA-col1 (layer 1, head 5; light orange).

Figure 4 illustrates the average *ESS* across all datasets for each combination of layer and column attention head. Notably, the *ESS* is more pronounced in low-gap, compared to high-gap datasets, but the positive-to-negative trend over layers and heads remains consistent. That trend suggests learning of attention heads in later layers of the MSA-Transformer pay more attention to sequences that are evolutionarily similar to one another, whereas layer one, head five, is more attentive to distant pairs, or (conversely) pays less attention to close homologs.

**Figure 4.**
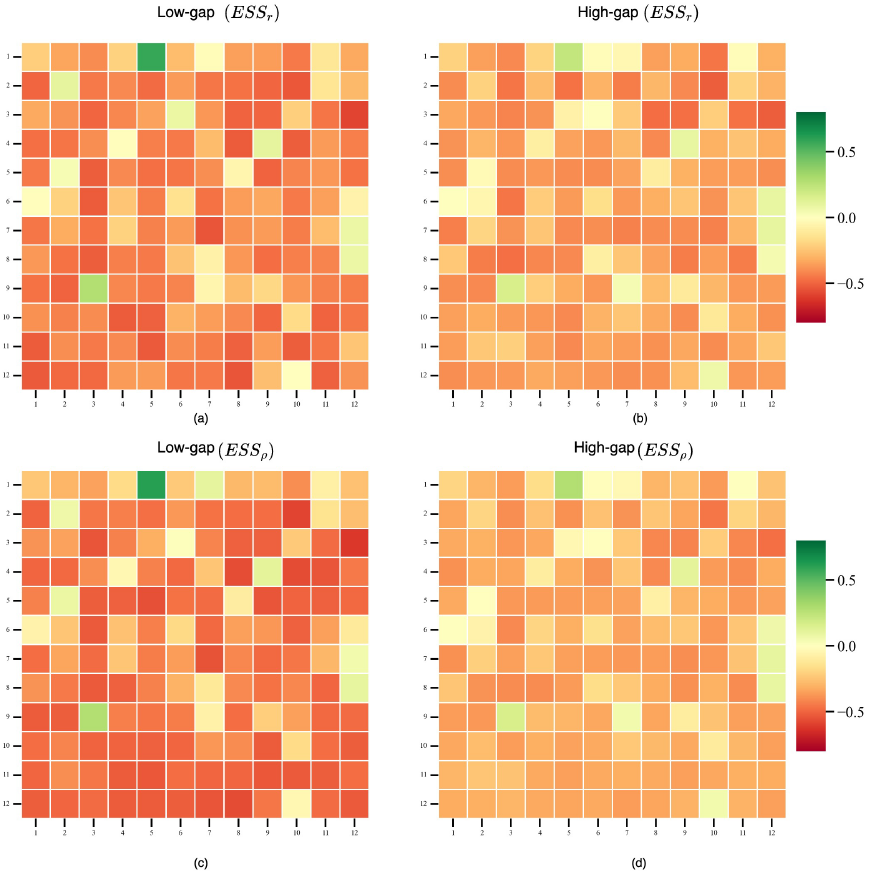
Average *ESS* for the LG matrix against all column attention heads of MSA-Transformer. x-axis show the different heads and y-axis captures the different layers of the MSA-Transformer. (a) Average *ESS*_*ρ*_ for low-gap datasets; (b) average *ESS*_*ρ*_ for high-gap datasets; (c) average *ESS*_*r*_ for low-gap datasets; (d) average *ESS*_*r*_ for high-gap datasets.

It remains unclear why, in certain datasets, the MSA-col1 exhibits only a low positive *ESS* while the MSA-col2 displays a strong negative *ESS*. This phenomenon is more pronounced in high-gap compared to low-gap ones. The average absolute difference in *ESS*_*r*_ between MSA-col2 and MSA-col1 is 0.28 for high-gap datasets (0.003 in low-gap), while the average absolute difference in *ESS*_*ρ*_ is 0.19 for high-gap datasets (0.004 in low-gap) (see Supplementary Table S3). A one-sided Wilcoxon rank-sum test confirms that the absolute *ESS* score (*ESS*_*ρ*_ and *ESS*_*r*_) between MSA-col2 and MSA-col1 is significantly greater in high-gap than in low-gap datasets.

### 3.4 pLMs capture relationships at different evolutionary time scales

We note that protein language models are trained on sequences from within and across diverse families. Thus, pLMs need to accommodate homology at widely different levels of similarity; the variable use of attention heads in the MSA-Transformer is suggestive of that range.

We hypothesised the ability to recover evolutionary relationships changes with the time sequences have evolved away from one another, and designed a series of tests to establish if pLMs operated differently at different evolutionary time scales.

#### 3.4.1 Both MSA-Transformer and ESM2 capture closely homologous groups, but gaps impede MSA-Transformer

We first assess the accuracy by which close homology is recovered by different pLM representations in terms of, and based on K-nearest neighbours. We vary the meaning of “close” by setting 5 ≤ K ≤ 20 to identify groups of sequences, as the closest by evolutionary distance from a reference sequence, in turn guided by an additive tree.

We measure the concordance of groups of *K* nearest sequences (in pLM embedding and tree space) by the Jaccard similarity coefficient (*LHS*; see Equation 3).Each sequence from the 100 Pfam datasets serves as a reference sequence once; 14 datasets are excluded as they cannot by processed by the MSA-Transformer. We bin groups identified by a reference sequence into four overlap categories: disagree (*LHS* = 0), low (0 *< LHS* ≤ 0.5), high (0.5 *< LHS <* 1), and agree (*LHS* = 1).

In general, the overlap is low across the board, suggesting that recovering the same nearest neighbours is challenging. In the low-gap category, MSA-col1 has the highest percentage of sequence groups that “agree” or are “high” in overlap outcomes for all values of *K* in both evolutionary distance and by its column attention (Figure 5).ESM2-embed performs similarly, but does so also for the high-gap category, suggesting that it is able to recover the same closely homologous sequences to a higher standard than other pLMs even in the presence of gaps.

**Figure 5.**
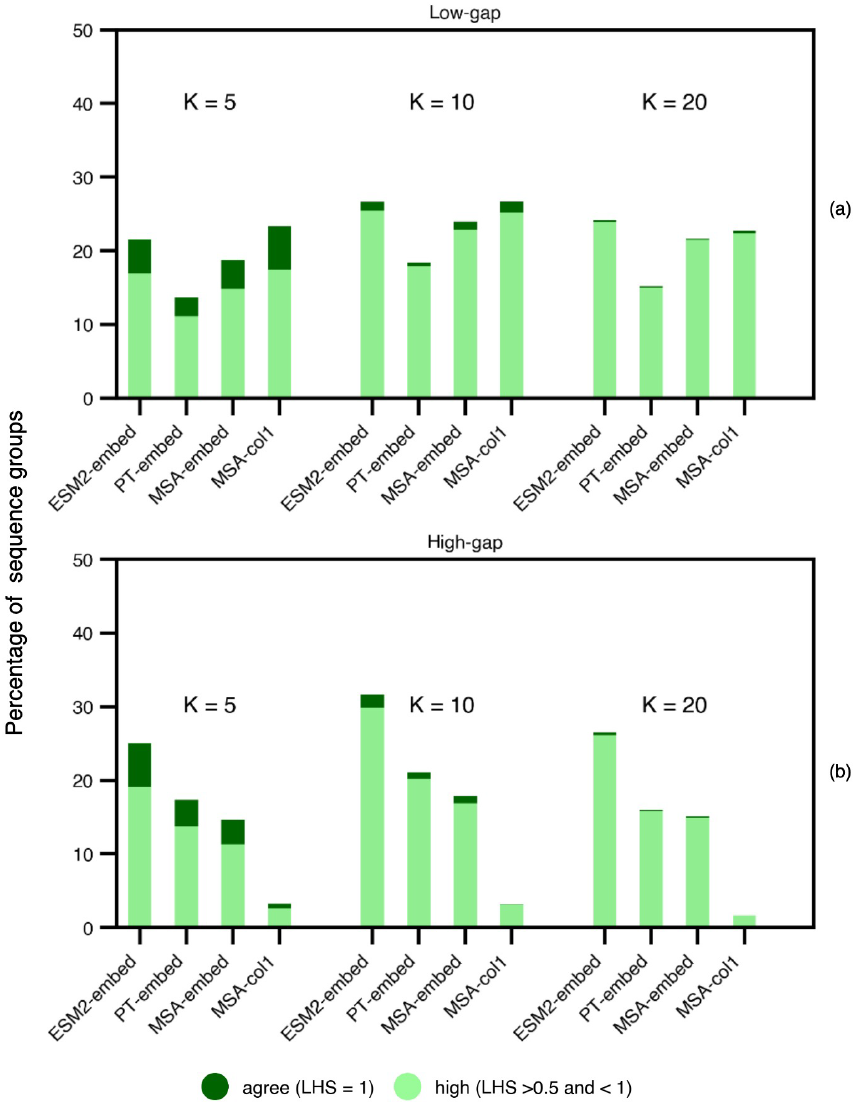
The percentage of close homologous sequence groups of size *K* drawn from all datasets, indicating different *LHS* overlap categories using the y-axis, and pLM embedding using the x-axis. We separate analyses for (a) low-gap and (b) high-gap datasets.

#### 3.4.2 Reproducing remote Le-Gascuel distances is easier for single-sequence pLMs

Here, we hypothesise that pLM representations primarily accommodate *broader* evolutionary relationships, as opposed to *finer* details that help distinguish close members of any particular protein family. Previously, we defined groups of sequences at the same high level of homology, referencing distance to investigate if pLM representations account for such groupings. To address pLMs’ sensitivity to remote homology, we here measure the concordance between *distances* in LG trees and *distances* in pLM representations, at different levels of “granularity”.

To probe different levels of granularity we next carefully selected subsets of sequence pairs indexed in **M**_LG_ vs **M**_pLM_ (Table 2).The trees in the Pfam datasets encompass a wide range of evolutionary time scales, so we first look at them individually, relative to a reference sequence. Specifically, we select the ten sequences that evenly sample all sorted evolutionary distances from the reference. We then retrieve the corresponding sequences in the pLM matrix to evaluate the extent distances in the embedding space are correlated; to this end, we define a “broad” *ESS* specific to a Pfam dataset as the average of correlation coefficients (for all possible reference sequences).

We repeat the above procedure to gauge whether pLMs are capable of recovering evolutionary distances at the finest scale applicable to each tree. To evaluate this, we define the “fine” *ESS* based on the ten *closest* sequences to each reference sequence within each dataset, and determine if their closeness in the LG matrix is correlated with the same in the embedding matrix for each pLM. We expected the correlation between the evolutionary distance and the similarity between the embeddings to be weaker, since all ten data points are closer to one another and therefore easier to mix up. We compare the “broad” and “fine” against “all” sequences for a reference sequence.

Generally and as expected, the “broad” *ESS*_*r*_ is higher than “all” *ESS*_*r*_, and the “fine” *ESS*_*r*_ is lower than the “all” *ESS*_*r*_. This trend is particularly pronounced in datasets where the pLMs achieve better *ESS*_*r*_ values compared to those with poorer *ESS*_*r*_ performance. Figure 6 shows analyses for the PT-embed and MSA-col1 pLM representations for all applicable datasets; ESM2-embed is trending similarly to PT-embed, while MSA-embed is concordant with MSA-col1 (see Supplementary Figure S6(c)). Similar trends are visible with *ESS*_*ρ*_ (see Supplementary Figure S6(a), Figure S6(b)).

**Figure 6.**
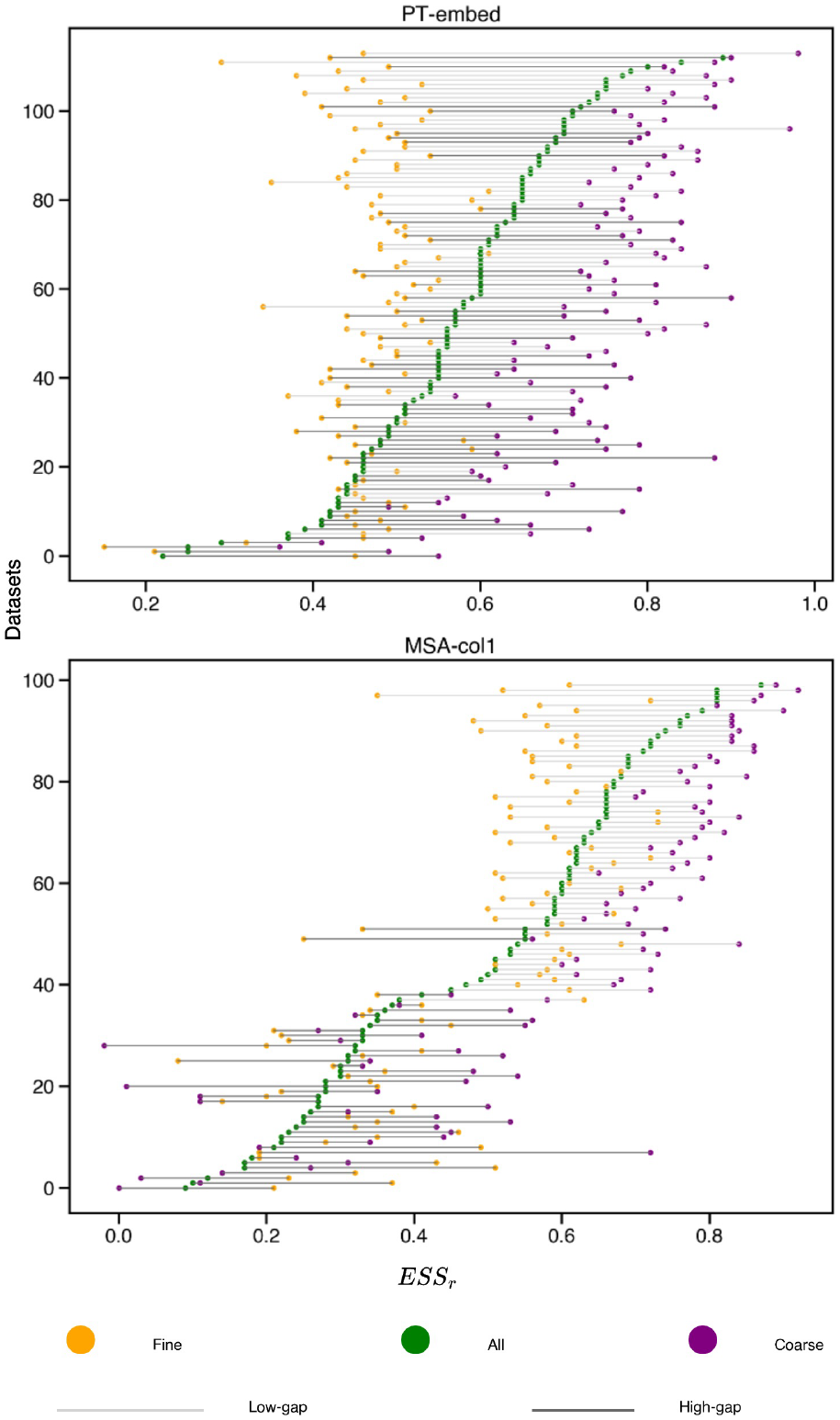
Dot plot representing the “fine” (yellow) and “broad” (purple) *ESS*_*r*_ (shown on x-axis), as well as the *ESS*_*r*_ including “all” (pairs of) sequences (green), for all Pfam datasets sorted on the y-axis in ascending order (top-to-bottom) by the “all” correlation. We include LG matrix vs. PT-embed (top panel) and LG matrix vs. MSA-col1 (bottom panel). Lines in light grey represent low-gap and dark grey represent high-gap Pfam datasets.

pLMs capture evolutionary relationships at the broad level at a consistently high level (Figure 6).This is reflected in the significant reductions in variance for “broad” *ESS*_*r*_: 20% for ESM2-embed and 27% for PT-embed. MSA-col1 and MSA-embed does not show such overall reductions but demonstrates a 35% and 11% reduction respectively for low-gap datasets. Interestingly, evolutionary relationships at the finer level are also reproduced consistently well across different families. In case of “fine” *ESS*_*r*_, the variance reduces by 43% for ESM2-embed, 72% for PT-embed, 40% for MSA-col1 and 37% for MSA-embed. However, the accurate reproduction of evolutionary distances across the full scale (as indicated by the “all” *ESS*_*r*_) varies greatly between different protein families.

To further illustrate if a pLM has a “sweet spot” for capturing evolutionary relationships in terms of reproducing their pairwise distances, we devise another test. As above, a reference sequence is grouped with 10 other members of the same tree, either because they are the closest or represent the range of distances in the tree. Regardless, by random chance, a greater *mean* distance in the group implies greater *variance*. We re-use the same groups, now aggregating them from *all* the trees, and stratifying them by their mean and variance. The resulting bins enable us to look at particulars of groups where members are *close homologs*, which is when the *mean* distance is small, or *remote homologs*, which is when the *mean* distance is large; we refer to the middle ground as *intermediate homologs*. Non-exclusive to the above labels, we distinguish groups where members are *highly divergent* which is when the variance of distance is large and implies a level of heterogeneity amongst members, or *not divergent*, which occurs when members are similar to one another; the middle ground here we refer to as *moderately divergent*.

Each pair of the above mean distance and variance of distance represent recurring phylogenetic tree-motifs that are useful to inspect in separation, e.g. dense clades of homologous sequences, with and without an outlier reference sequence, or sparse trees, with very divergent sequences. We illustrate a couple of these motifs in Figure 7,and show *ESS*_*r*_ (*y*-axis in the 2D density plots) for the stratified data vis-á-vis their mean in distance (*x*-axis), for each of the pLMs. We note the *same* groups are analysed for each pLM, so the variance of distance is identical for each label-pair, though some label-pairs are well represented and some are much less so. The key metric of success is to achieve an *ESS*_*r*_ of 1, which implies that the distance from LG is reproduced.

**Figure 7.**
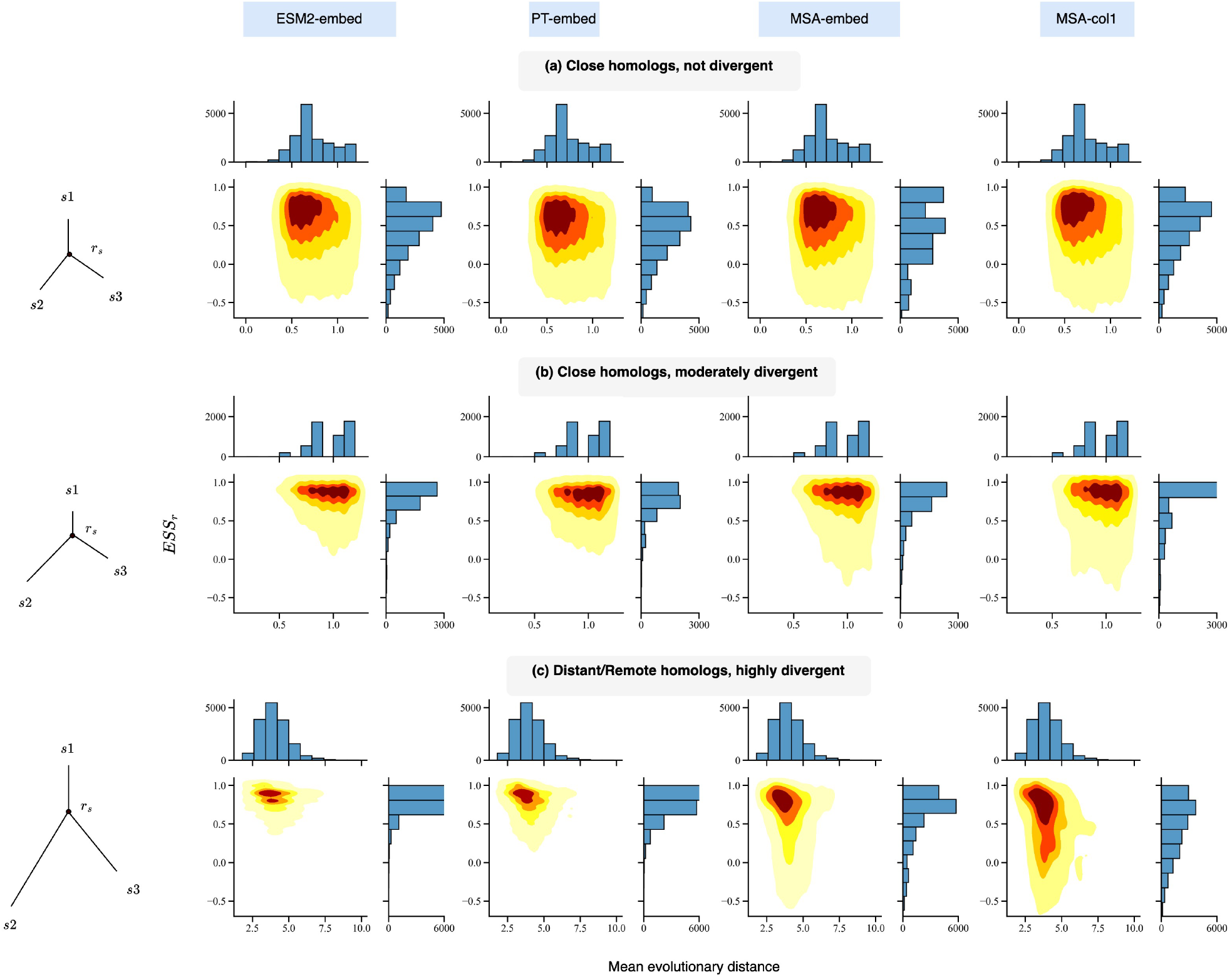
Analysis of *ESS*_*r*_ for different evolutionary distances for all pLM representations. Density of *ESS*_*r*_ for “close homologs, not divergent” (a), “close homologs, moderately divergent” (b) and “distant/remote homologs, highly divergent” (c) relative to mean evolutionary distance in each group (from reference sequence). Left-hand side plots in each group show the caricatures of nominal but common phylogenetic tree-motifs (*r*_*s*_ is a reference sequence and *s*_1_, *s*_2_, *s*_3_ are members of its group).

When the sequence pairs are highly divergent, ESM2-embed is closest (of pLMs) to reproducing the classical LG-based relationship, in particular at longer evolutionary distances (see Figure 7(c) where distance [*x*] ranges 2-6, but also Figure 7(b) where distance is closer to 1). We note that in the case of moderately and highly divergent groups, the distribution of *ESS*_*r*_ for ESM2 is significantly greater than that of all pLM representations. For close, not divergent groups, ESM2 still has a significantly greater *ESS*_*r*_, except for MSA-col1 which is statistically indistinguishable (see Figure 7(b) where distance is less than 1 and members form a dense, non-divergent group). Between the two MSA-Transformer representations, MSA-col1 is significantly closer to LG-based distances in close and not divergent groups, but this order is reversed in moderately and highly divergent groups.

### 3.5 Less than 10% of the neurons capture sequence based phylogenetic evolutionary signal between homologous proteins

Here we set out to understand how sequence-based phylogenetic signal is organised in the embedding space of a pLM. We conduct this probe for each “gap” category and for each pLM representation using the top 5-performing Pfam datasets (in terms of *ESS*_*r*_) (see Section 2.2.3).Supplementary Figure S7(a) shows the datasets in the form of the Venn diagram with some shared, and some exclusive to any given pLM.

This “neuron saliency” probe relies on training an Elastic Net regressor [31] to identify neurons (in the output dimension) that best predict the evolutionary distance between any pair of homologous protein sequences. Each pair is represented by the *absolute difference* at selected neurons. To select the neurons that will contribute to training, we only use those with an activation variance greater than the 20th percentile of the aggregate values for a particular dataset, a selection process that is independent of the evolutionary distance. We randomly sample 9000 instances of two sequences within a dataset and apply a train/test split ratio of 70/30. We assess the trained model’s performance on the test set based on the *R*^2^ score against the evolutionary distances. For each dataset, we repeat the steps ten times using a different random sample.

In agreement with previous, zero-shot analyses, the average *R*^2^ between the Elastic Net prediction for a sequence and the actual evolutionary distance is greater in low-gap than in high-gap datasets, across all pLM representations. Moreover, *R*^2^ varies less in low-gap than in high-gap category (see Supplementary Figure S7(b)).

The coefficients at inputs to the trained Elastic Net represent the corresponding neuron’s *saliency* to predicting the evolutionary distance. These coefficients are analysed in aggregate by summing their absolute values across ten runs. Subsequently, we bin neurons into the top 10%, 25%, 50%, 75% and bottom 25% based on these sums, arranging them in descending order.

In single-sequence pLMs, the phylogenetic signal is concentrated to a greater number of neurons for high-gap datasets compared to low-gap datasets; this is unlike the MSA Transformer that specifically embeds a sequence in context of an alignment, and is therefore not directly comparable. In the case of ESM2-embed, the number of salient neurons is approximately three times that of the other pLM representations. This may be due to the larger embedding dimension of ESM2, which could also give the ESM2 model an advantage with high-gap datasets, as indicated by the high *R*^2^ score distribution in Supplementary Figure S7(b). Figure 8(a) shows the distribution of the neurons in the top 10% bin for each pLM representation.

**Figure 8.**
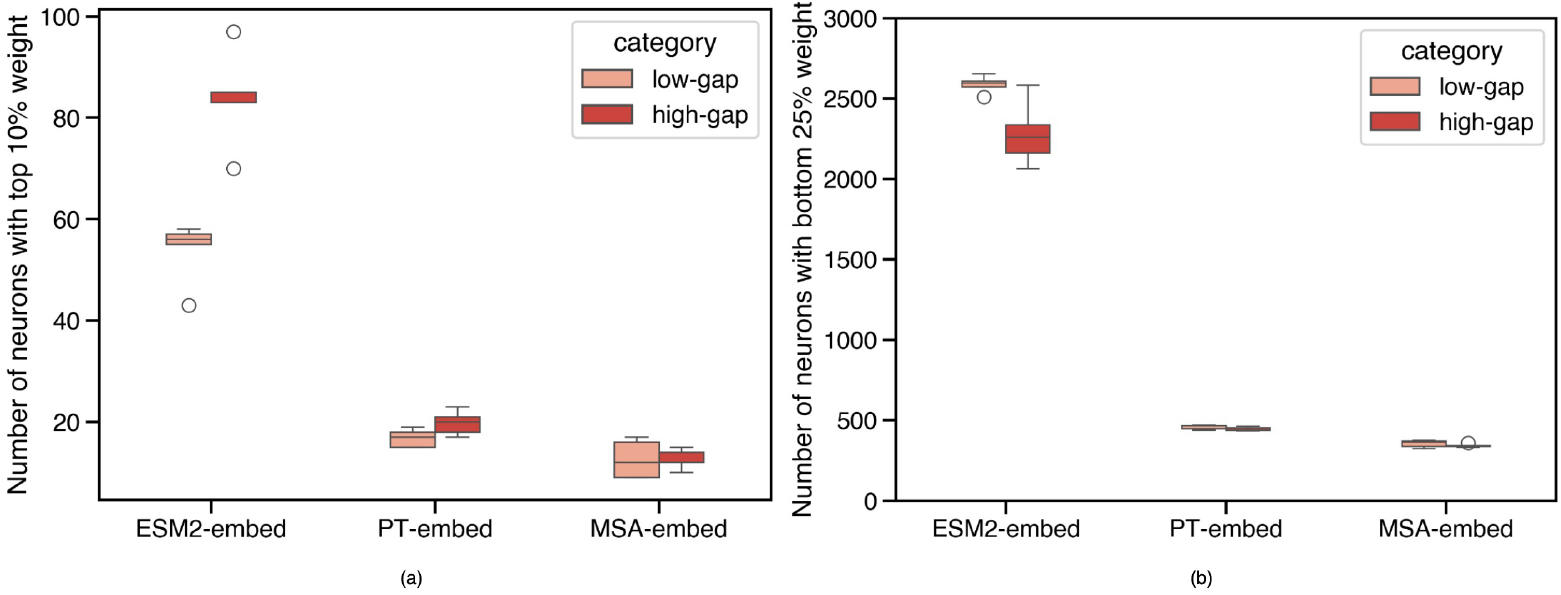
Probing the saliency of neurons. (a) Distribution of the top salient neurons identified by elastic net regression for each pLM representation. ESM2 model shows a high number of neurons used for capturing evolutionary distances between protein sequences. (b) Distribution of the bottom neurons identified by elastic net regression for each pLM representation.

Next, we evaluate our scoring strategy through so-called neuron ablations [31].In this process, we perform a similar analysis as highlighted in section 3.2,but focus solely on the top and bottom neurons identified for each dataset and masking out the rest. We observe that the *ESS*_*r*_ of bottom neurons is lower compared to the *ESS*_*r*_ reported by the top-saliency neurons (see Supplementary Tables S4-S6), thus validating our scoring strategy.

We observe an increase in the *ESS*_*r*_ for the top-saliency neurons, with the score decreasing as more neurons were included. For ESM2-embed and MSA-embed, the peak *ESS* typically occurs when only the top 10% of neurons are used, with a few exceptions. In contrast, for PT-embed, the *ESS*_*r*_ peaks when the top 25% of neurons are considered in most datasets. This suggests that the evolutionary relationships between homologous proteins are localised rather than distributed throughout the embedding space. These localised neurons may be specialised in capturing the unique characteristics of the sequences in that dataset, making the representation more tailored and less generalised. Supplementary Tables S4-S6 show the *ESS*_*r*_ for all, top and bottom neurons for the datasets used for training elastic net.

The neuron saliency probe gives us some insight into the organisation of the embedding space of each pLM. For each pLM, it is noteworthy that the combined top 10% of neurons are unique for each dataset, indicating a global family-wise organization of the embedding space. There are some overlapping neurons (see Figure 9) shared among datasets but none that are shared across all ten datasets considered for this analysis. The overlapping neurons hint at the polysemantic properties of neurons in the neural networks [32], where features are distributed across neurons and one neuron can encode multiple features (i.e. they activate in multiple semantic distinct context).

**Figure 9.**
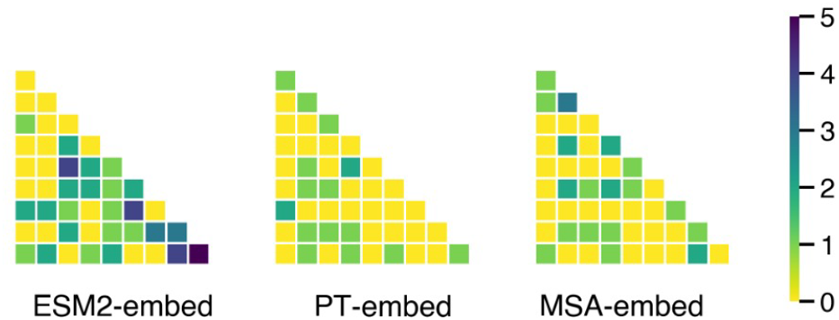
Heatmap showing the number of superimposed neurons in the top 10% of neurons for datasets used in evolutionary probe. x-axis and y-axis represent the different datasets used for evolutionary probe. ESM2-embed show a high overlap of neurons compared to the other pLM representation; this might be a characteristic of the protein datasets used for probe rather than property of that particular pLM.

### 3.6 ESM2 agrees more with evolutionary models than ESM3

Structure is often considered as more evolutionarily conserved than sequence [33].With the advent of multimodal protein language models (pLMs) such as ESM3 and ProstT5, we hypothesise that these multimodal models outperform sequence-based pLMs in both the *ESS* and *LHS* metrics. For this analysis, we utilise the mean embeddings (see Section 2.1.2)from the output layers of ESM3 (referred to as ESM3-embed) and ProstT5 (referred to as PST-embed).

For the *LHS* analysis, our results show that ESM2-embed consistently has the highest percentage of sequence groups that “agree” or exhibit “high” overlap outcomes across all values of *K*. Additionally, PST-embed outperforms PT-embed (refer to Tables 4 and 3).

**Table 3.** *LHS* for low-gap

| Name | k | high | agree |
| --- | --- | --- | --- |
| ESM2-embed / ESM3-embed | 5 | **16.90** / 6.05 | **4.61** / 1.02 |
| PT-embed / PST-embed | 5 | 11.12 / **13.05** | 2.54 / **2.89** |
| ESM2-embed / ESM3-embed | 10 | **25.46** / 8.83 | **1.18** / 0.17 |
| PT-embed / PST-embed | 10 | 17.86 / **19.20** | 0.54 / **0.56** |
| ESM2-embed / ESM3-embed | 20 | **23.93** / 6.89 | **0.25** / 0.00 |
| PT-embed / PST-embed | 20 | 15.01 / **16.78** | 0.12 / **0.16** |

**Table 4.** *LHS* for high-gap

| Name | k | high | agree |
| --- | --- | --- | --- |
| ESM2-embed / ESM3-embed | 5 | **19.12** / 8.39 | **5.95** / 1.44 |
| PT-embed / PST-embed | 5 | 13.77 / **14.62** | 3.55 / **4.11** |
| ESM2-embed / ESM3-embed | 10 | **29.86** / 11.37 | **1.77** / 0.30 |
| PT-embed / PST-embed | 10 | 20.19 / **22.51** | 0.91 / **1.05** |
| ESM2-embed / ESM3-embed | **20** | **26.11** / 7.76 | **0.40** / 0.07 |
| PT-embed / PST-embed | 20 | 15.88 / **19.06** | 0.08 / **0.26** |

When comparing *ESS* between PT-embed and PST-embed, there is no significant difference in *ESS*_*r*_. However, PST-embed has a slightly higher *ESS*_*ρ*_ (Wilcoxon signed-rank test, *p <* 0.05) across both low-gap and high-gap datasets. In contrast, both *ESS* metrics for ESM2-embed are significantly higher than those for ESM3-embed across all gap categories (see Supplementary Figure S8).

Finally, when evaluating the “sweet spot” for capturing evolutionary relationships (as in Section 3.4.2), ESM2-embed outperforms ESM3-embed significantly across all three bins (close not divergent homologs, close moderately divergent homologs, and distant/remote highly divergent homologs (see Figure 7)).For PST-embed and PT-embed there is no statistically significant difference between them, except in the case of close and moderately divergent homologs, where PT-embed is significantly greater (see Supplementary Figure S9).

## 4 Discussion

Our study explores the overlap between classical phylogenetic methods and pLMs in capturing evolutionary information from homologous protein sequences. The pLMs chosen for analysis share many similar characteristics while also possessing distinctive features that make them particularly interesting to compare. ESM2 and ProtT5 are both trained on UniRef50 sequences and have a comparable number of parameters (15 billion vs. 11 billion), yet they feature different architectures, training objectives and embedding dimensions (5120 vs. 1024). In contrast, the alignment-trained model MSA-Transformer is relatively small, with 12 layers and 100 million parameters, featuring a 768-dimensional embedding. Each pLM can produce a numerical vector representation for a given protein sequence, either with or without an MSA. These numerical representations are meaningful when processed or compared against each other, providing insights into the internal workings of the pLMs as well the information captured by the representations themselves. They also enable cross-model comparison and analysis.

Deep learning models are frequently characterised as “black boxes” due to the difficulty in deciphering their decision-making processes. By employing a range of probes or stimuli, we can illuminate the internal workings and learning patterns of these models. In this paper, we utilise various probes, ranging from straightforward to intricate, to enhance our understanding of pLMs. The simplest one-hot correlation test reveals the first 30% of the layers in the single-sequence model are not context-aware. This is consistent with the way deep learning models learn patterns in a bottom-up fashion, starting with simple features in early layers and progressively capturing more complex patterns in deeper layers. In contrast, the MSA-based model, which uses both row and column attention mechanisms, shows a different learning pattern compared to single-sequence models. This is evident from the increasing correlation between pLM representations and one-hot encoding in the later layers of the MSA-based model, whereas in single-sequence models, this correlation typically decreases from high to low.

When we compare the distances between pLM sequence representations and the tree distances in phylogenetic trees for homologous protein sequences, pLM performance varies between low- and high-gap datasets, with MSA-based representations under-performing in high-gap settings. High-gap MSAs, which reflect more complex relationships among individual sequences, might benefit from models with greater parameter capacity or learning ability to capture these intricate patterns. This is evident as single-sequence models show improved alignment with evolutionary models for high-gap datasets, even without relying on the MSA. Conversely, for low-gap MSAs, where sequences are more conserved, the MSA-Transformer exhibits more consistent performance (i.e., no dataset has *ESS*_*ρ*_ or *ESS*_*r*_ ≤ 0.3). In these cases, sequence alignment data proves advantageous for elucidating relationships between highly similar sequences. Incorporating MSA information has been shown to enhance the performance of AlphaFold [34]by providing additional evolutionary context that aids in accurate structure prediction. Simi-larly, single-sequence models might benefit from incorporating MSA information for highly conserved proteins into their training routines, potentially enhancing their ability to capture the evolutionary relationships of both highly conserved and non-conserved protein sequences.

Although ProtT5 and ESM2 are the largest models in their respective categories, with substantial learning capacities, ESM2 has an output dimension that is five times larger than that of ProtT5. This difference might explain why relationships based on ESM2 shows better overall agreement with the evolutionary model. Additionally, ESM2 also achieves the best performance in matching the k-nearest neighbors with the phylogenetic tree and reproducing remote Le-Gascuel distances. This demonstrates that large dimension size enhances the model’s representational capacity, allowing it to capture more nuanced relationships between sequences even in the absence of MSA information.

Finally, we assessed sequence representations from models trained with structural and functional information and obtained mixed results. ProstT5 embeddings either outperformed or performed similarly to those from ProtT5-XXL UniRef50, while embeddings from ESM3 did not surpass those from ESM2. This suggests that the larger hidden dimension in ESM2 outweighs the potential benefits of structural information when encoding phylogenetic relationships based on protein sequences.

In this study, we have explored multiple approaches to investigate how homologous relationships are captured in pLMs. However, our study is limited as it is based solely on protein sequences. Future research could benefit from extending this methodology to incorporate additional molecular sequence characteristics, potentially providing a more comprehensive understanding of these relationships.

## References

[1] B. Q. Minh, H. A. Schmidt, O. Chernomor, D. Schrempf et al., “IQ-TREE 2: New Models and Efficient Methods for Phylogenetic Inference in the Genomic Era,” Molecular Biology and Evolution, vol. 37, no. 5, pp. 1530–1534, 02 2020. [Online]. Available: 10.1093/molbev/msaa015

[2] P. S. D. Morgan N. Price and A. P. Arkin, “Fasttree 2 – approximately maximum-likelihood trees for large alignments,” PLOS ONE, vol. 5, no. 3, pp. 1–10, 03 2010. [Online]. Available: 10.1371/journal.pone.0009490

[3] S. Q. Le and O. Gascuel, “An Improved General Amino Acid Replacement Matrix,” Molecular Biology and Evolution, vol. 25, no. 7, pp. 1307–1320, 03 2008. [Online]. Available: 10.1093/molbev/msn067

[4] Q. Zhan, Y. Ye, T.-W. Lam, S.-M. Yiu et al., “Improving multiple sequence alignment by using better guide trees,” BMC Bioinformatics, vol. 16, no. 5, p. S4, Mar. 2015.

[5] A. Tóth, A. Hausknecht, I. Krisai-Greilhuber, T. Papp et al., “Iteratively refined guide trees help improving alignment and phylogenetic inference in the mushroom family bolbitiaceae,” PLOS ONE, vol. 8, no. 2, pp. 1–14, 02 2013. [Online]. Available: 10.1371/journal.pone.0056143

[6] D. Brown and K. Sjölander, “Functional classification using phylogenomic inference,” PLoS Comput Biol, vol. 2, no. 6, p. e77, Jun. 2006.

[7] Z. Jiang, “Protein function predictions based on the phylogenetic profile method,” Crit Rev Biotechnol, vol. 28, no. 4, pp. 233–238, 2008.

[8] M. Musil, R. T. Khan, A. Beier, J. Stourac et al., “FireProtASR: A Web Server for Fully Automated Ancestral Sequence Reconstruction,” Briefings in Bioinformatics, vol. 22, no. 4, p. bbaa337, 12 2020. [Online]. Available: 10.1093/bib/bbaa337

[9] G. Foley, A. Mora, C. M. Ross, S. Bottoms et al., “Engineering indel and substitution variants of diverse and ancient enzymes using graphical representation of ancestral sequence predictions,” PLOS Computational Biology, vol. 18, no. 10, pp. 1–35, 2022. [Online]. Available: 10.1371/journal.pcbi.1010633

[10] A. Rives, J. Meier, T. Sercu, S. Goyal et al., “Biological structure and function emerge from scaling unsupervised learning to 250 million protein sequences,” Proceedings of the National Academy of Sciences, vol. 118, no. 15, p. e2016239118, 2021. [Online]. Available: https://www.pnas.org/doi/abs/10.1073/pnas.2016239118

[11] K. Kaminski, J. Ludwiczak, K. Pawlicki, V. Alva et al., “pLM-BLAST: distant homology detection based on direct comparison of sequence representations from protein language models,” Bioinformatics, vol. 39, no. 10, p. btad579, 09 2023. [Online]. Available: 10.1093/bioinformatics/btad579

[12] B. Hie, E. D. Zhong, B. Berger, and B. Bryson, “Learning the language of viral evolution and escape,” Science, vol. 371, no. 6526, pp. 284–288, Jan. 2021.

[13] B. L. Hie, K. K. Yang, and P. S. Kim, “Evolutionary velocity with protein language models predicts evolutionary dynamics of diverse proteins,” Cell Systems, vol. 13, no. 4, pp. 274–285.e6, 2022. [Online]. Available: https://www.sciencedirect.com/science/article/pii/S2405471222000382

[14] B. G. Iovino and Y. Ye, “Protein embedding based alignment,” BMC Bioinformatics, vol. 25, no. 1, p. 85, Feb 2024. [Online]. Available: 10.1186/s12859-024-05699-5

[15] L. Pantolini, G. Studer, J. Pereira, J. Durairaj et al., “Embedding-based alignment: combining protein language models with dynamic programming alignment to detect structural similarities in the twilight-zone,” Bioinformatics, vol. 40, no. 1, p. btad786, 01 2024. [Online]. Available: 10.1093/bioinformatics/btad786

[16] J. Meier, R. Rao, R. Verkuil et al., “Language models enable zero-shot prediction of the effects of mutations on protein function,” bioRxiv, 2021. [Online]. Available: https://www.biorxiv.org/content/early/2021/11/17/2021.07.09.450648

[17] Y. Qiu, J. Hu, and G.-W. Wei, “Cluster learning-assisted directed evolution,” Nat Comput Sci, vol. 1, no. 12, pp. 809–818, Dec. 2021.

[18] B. J. Wittmann, Y. Yue, and F. H. Arnold, “Informed training set design enables efficient machine learning-assisted directed protein evolution,” Cell Syst, vol. 12, no. 11, pp. 1026–1045.e7, Aug. 2021.

[19] D. S. Umberto Lupo and A.-F. Bitbol, “Protein language models trained on multiple sequence alignments learn phylogenetic relationships,” Nature Communications, vol. 13, no. 1, p. 6298, Oct. 2022.

[20] R. V. J. M. Roshan M Roa, Jason Liu et al., “Msa transformer,” in Proceedings of the 38th International Conference on Machine Learning, ser. Proceedings of Machine Learning Research, M. Meila and T. Zhang, Eds., vol. 139. PMLR, 18–24 Jul 2021, pp. 8844–8856. [Online]. Available: https://proceedings.mlr.press/v139/rao21a.html

[21] Z. Lin, H. Akin, R. Rao, B. Hie et al., “Evolutionary-scale prediction of atomic-level protein structure with a language model,” Science, vol. 379, no. 6637, pp. 1123–1130, 2023. [Online]. Available: https://www.science.org/doi/abs/10.1126/science.ade2574

[22] A. Elnaggar, M. Heinzinger, C. Dallago, G. Rehawi et al., “Prottrans: Toward understanding the language of life through self-supervised learning,” IEEE Transactions on Pattern Analysis and Machine Intelligence, vol. 44, no. 10, pp. 7112–7127, 2022.

[23] T. Hayes, R. Rao, H. Akin, N. J. Sofroniew et al., “Simulating 500 million years of evolution with a language model,” bioRxiv, 2024. [Online]. Available: https://www.biorxiv.org/content/early/2024/07/02/2024.07.01.600583

[24] M. Heinzinger, K. Weissenow, J. G. Sanchez, A. Henkel et al., “Prostt5: Bilingual language model for protein sequence and structure,” bioRxiv, 2023. [Online]. Available: https://www.biorxiv.org/content/early/2023/07/25/2023.07.23.550085

[25] L. W. M. Q. Jaina Mistry, Sara Chuguransky et al., “Pfam: The protein families database in 2021,” Nucleic Acids Research, vol. 49, no. D1, pp. D412–D419, 10 2020. [Online]. Available: 10.1093/nar/gkaa913

[26] N. Kriegeskorte, M. Mur, and P. Bandettini, “Representational similarity analysis - connecting the branches of systems neuroscience,” Frontiers in Systems Neuroscience, vol. 2, 2008. [Online]. Available: https://www.frontiersin.org/articles/10.3389/neuro.06.004.2008

[27] S. Abnar, L. Beinborn, R. Choenni, and W. H. Zuidema, “Blackbox meets blackbox: Representational similarity and stability analysis of neural language models and brains,” CoRR, vol. abs/1906.01539, 2019. [Online]. Available: http://arxiv.org/abs/1906.01539

[28] F. S. Jaime Huerta-Cepas and P. Bork, “ETE 3: Reconstruction, analysis, and visualization of phylogenomic data,” Mol. Biol. Evol., vol. 33, no. 6, pp. 1635–1638, Jun. 2016.

[29] L. Valeriani, F. Cuturello, A. Ansuini, and A. Cazzaniga, “The geometry of hidden representations of protein language models,” bioRxiv, 2022. [Online]. Available: https://www.biorxiv.org/content/early/2022/10/26/2022.10.24.513504

[30] J. L. Hodges, “The significance probability of the smirnov two-sample test,” Arkiv för Matematik, vol. 3, no. 5, pp. 469–486, 1958. [Online]. Available: 10.1007/BF02589501

[31] F. Dalvi, N. Durrani, H. Sajjad, Y. Belinkov, A. Bau, and J. Glass, “What is one grain of sand in the desert? analyzing individual neurons in deep nlp models,” 2018. [Online]. Available: https://arxiv.org/abs/1812.09355

[32] N. Elhage, T. Hume, C. Olsson, N. Schiefer et al., “Toy models of superposition,” 2022. [Online]. Available: https://arxiv.org/abs/2209.10652

[33] K. Illergård, D. H. Ardell, and A. Elofsson, “Structure is three to ten times more conserved than sequence–a study of structural response in protein cores,” Proteins, vol. 77, no. 3, pp. 499–508, Nov. 2009.

[34] J. Jumper, R. Evans, A. Pritzel et al., “Highly accurate protein structure prediction with alphafold,” Nature, vol. 596, no. 7873, pp. 583–589, Aug 2021. [Online]. Available: 10.1038/s41586-021-03819-2

