## Supplementary Material for "Do Protein Language Models Learn Phylogeny?"

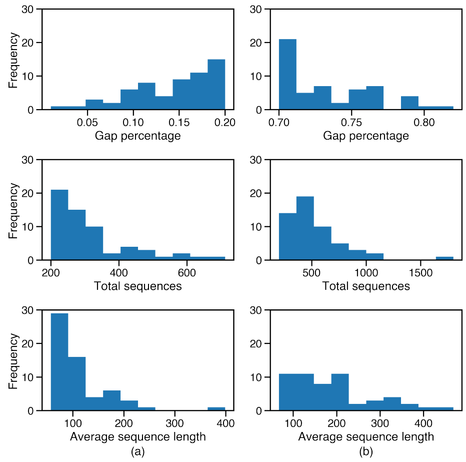


Fig. S1: Histogram plot showing the distribution of gap percentage, number and lengths of sequences in MSAs for low-gap (a) and high-gap (b) datasets.

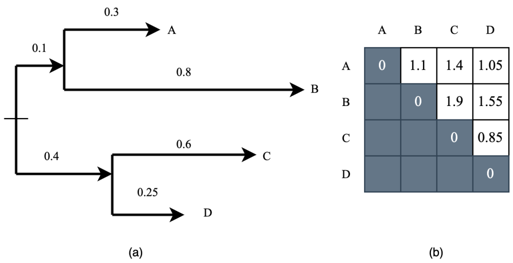


Fig. S2: Figure shows an example phylogenetic tree with four sequences - A, B, C and D (1) and corresponding LG matrix (2).

Table S1: Pfam datasets used for analysis.

| **Family low-gap** | **Domain low-gap** | **Family high-gap** | **Domain high-gap** |
|---|---|---|---|
| PF04298 | PF00158 | PF01055 | PF04205 |
| PF07907 | PF09278 | PF12686 | PF04427 |
| PF02702 | PF12002 | PF20060 | PF13508 |
| PF04961 | PF12775 | PF03914 | PF04264 |
| PF04461 | PF17941 | PF02586 | PF05192 |
| PF04403 | PF16658 | PF02365 | PF01156 |
| PF01668 | PF01706 | PF01139 | PF11799 |
| PF01450 | PF02261 | PF00617 | PF03796 |
| PF04304 | PF14278 | PF03150 | PF01288 |
| PF03975 | PF09361 | PF01297 | PF09994 |
| PF05635 | PF18565 | PF20415 | PF13280 |
| PF19609 | PF17146 | PF00856 | PF16124 |
| PF03862 | PF01948 | PF00902 | PF13356 |
| PF01219 | PF00707 | PF02104 | PF17852 |
| PF06421 | PF06071 | PF02811 | PF12804 |
| PF14842 | PF10437 | PF12276 | PF01510 |
| PF03788 | PF13667 | PF01368 | PF04122 |
| PF00934 | PF00189 | PF02383 | PF01369 |
| PF02049 | PF11760 | PF04051 | PF01302 |
| PF02617 | PF02650 | PF12704 | PF01266 |
| PF04341 | PF10369 | PF02517 | PF00006 |
| PF01052 | PF01037 | PF01728 | PF12697 |
| PF02700 | PF06130 | PF01885 | PF00557 |
| PF14841 | PF02594 | PF00636 | PF01388 |
| PF00986 | PF07554 | PF01196 | PF05257 |
| PF08439 | PF09269 | PF13423 | PF05226 |
| PF02805 | PF00366 | PF01926 | PF01149 |
| PF01313 | PF03719 | | |
| PF00831 | PF03880 | | |
| PF02073 | PF20554 | | |

Table S2: Total under performing datasets (where $ESS_\rho \leq 0.3$, $ESS_r \leq 0.3$ )

| **Name** | **low-gap** | **high-gap** |
|---|---|---|
| ESM2-embed | 5, 5 | 7, 3 |
| PT-embed | 1, 1 | 7, 5 |
| MSA-embed | 0, 0 | 28, 28 |
| MSA-col1 | 1, 1 | 38, 46 |

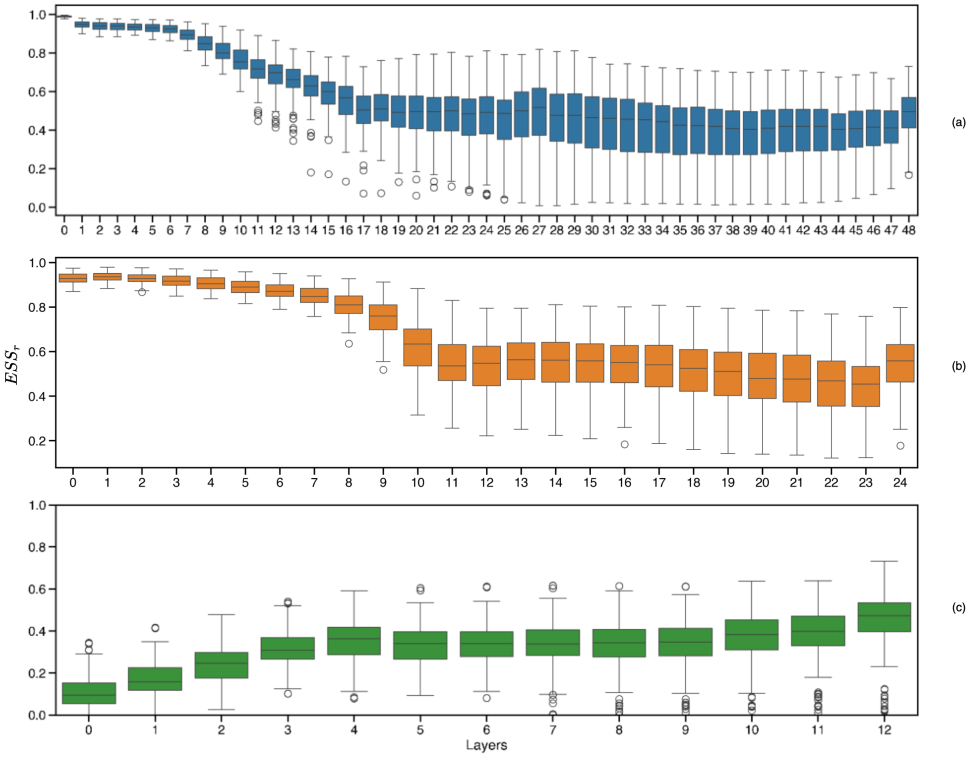


Fig. S3: Evolutionary similarity score ($ESS_r$; y-axis) between layer-specific pLM matrix (x-axis) and onehot matrix. Boxplots are based on 114 Pfam datasets for ESM2-embed (a) and PT-embed (b). In case of MSA-Transformer (c), boxplots are based on 100 Pfam entries.

Table S3: Difference in the absolute $ESS$ between MSA-col2 and MSA-col1. Mean and standard deviation are shown.

| **Dataset category** | $ESS_r$ | $ESS_\rho$ |
|---|---|---|
| lowgap | 0.003 (0.16) | 0.004 (0.12) |
| highgap | 0.28 (0.19) | 0.19 (0.17) |

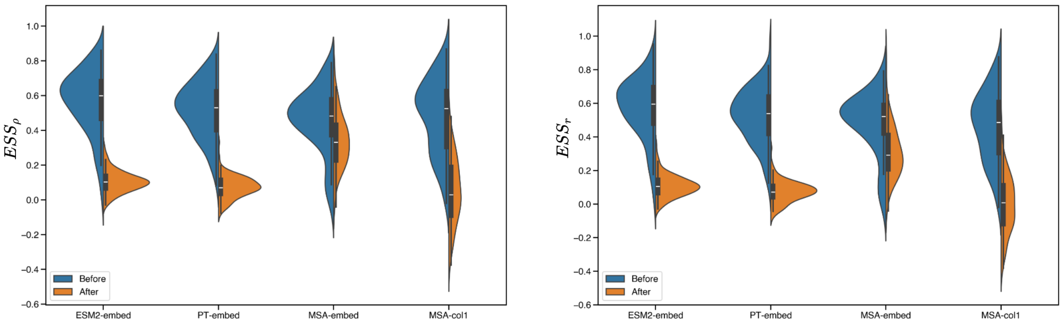


(a) (b)

Fig. S4: Distributions of $ESS_\rho$ (a) and $ESS_r$ (b) before and after shuffling of amino acids in the sequences. Sequence are shuffled for 80% of the total non-aligned sequences for ESM2-embed and PT-embed. For MSA-embed and MSA-col1, we shuffle on aligned sequences.

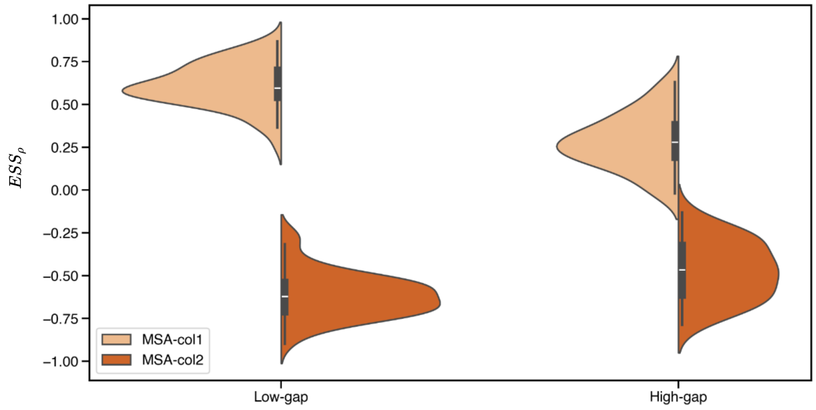


Fig. S5: Distributions of $ESS_\rho$ of select column attention heads in MSA-Transformer, across all applicable low-gap and high-gap datasets; the scores for MSA-col2 (layer three, head 12; dark orange) negatively mirrors MSA-col1 (layer 1, head 5; light orange).

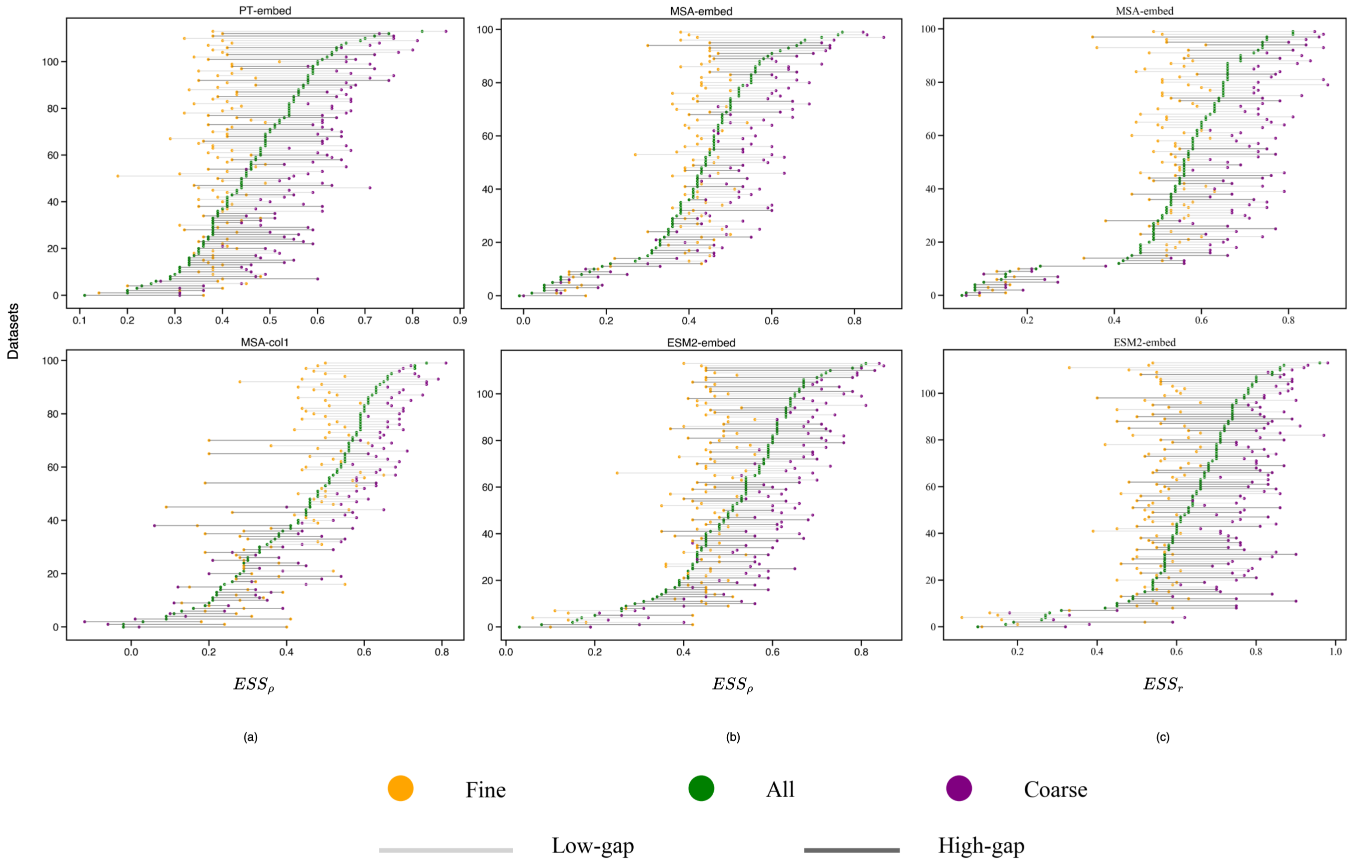


Fig. S6: Dot plot representing the "fine" (yellow) and "broad" (purple) $ESS_\rho$ (a, b) and $ESS_r$ (c) (shown on x-axis), as well as the $ESS$ including "all" (pairs of) sequences (green), for all Pfam datasets sorted on the y-axis in ascending order (top-to-bottom) by the "all" correlation. (a) We include LG matrix vs. PT-embed (top panel) and LG matrix vs. MSA-col1 (bottom panel). (b) We include LG matrix vs. MSA-embed (top panel) and ESM2 matrix vs. MSA-col1 (bottom panel). (c) We include LG matrix vs. MSA-embed (top panel) and LG matrix vs. ESM2-embed (bottom panel). Lines in light grey represent low-gap and dark grey represent high-gap Pfam datasets.

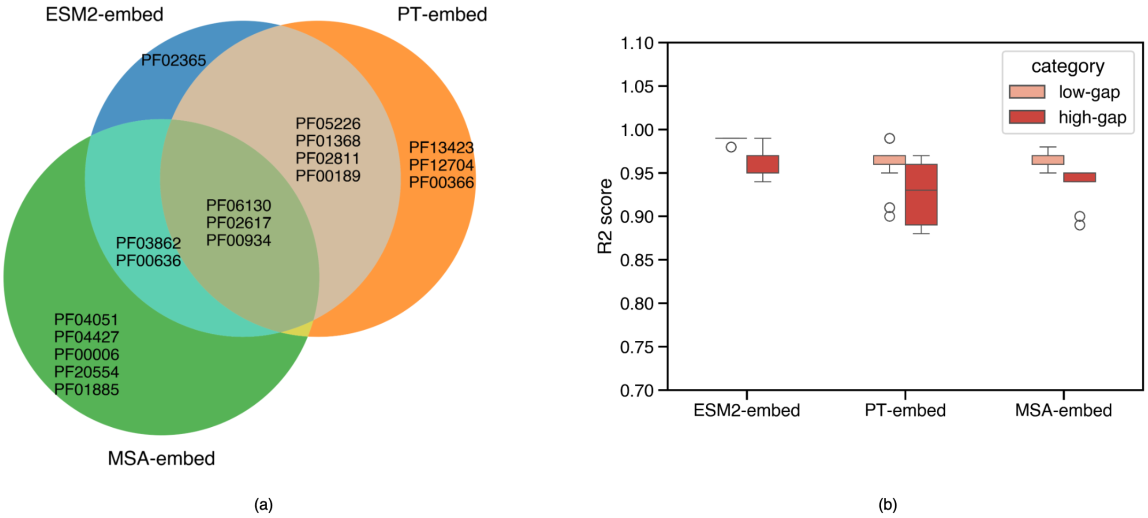


Fig. S7: A) Venn diagram showing datasets used to train elastic net regression for each pLM representation. B) Boxplot showing the distribution of $R^2$ score on test sets for each pLM representation. The score distibution is shown for both "gap" categories. .

Table S4: $ESS_r$ using salient and bottom neurons for ESM2-embed

| **Dataset** | **All** | **Top** | | | | **Bottom** |
|---|---|---|---|---|---|---|
| | | **(10%)** | **(25%)** | **(50%)** | **(75%)** | **(25%)** |
| PF03862 | 0.84 | 0.95 | 0.95 | 0.93 | 0.9 | 0.81 |
| PF00189 | 0.86 | 0.94 | 0.94 | 0.92 | 0.89 | 0.83 |
| PF02617 | 0.87 | 0.96 | 0.96 | 0.93 | 0.9 | 0.85 |
| PF00934 | 0.89 | 0.91 | 0.92 | 0.92 | 0.9 | 0.88 |
| PF06130 | 0.95 | 0.98 | 0.98 | 0.98 | 0.97 | 0.94 |
| PF02365 | 0.73 | 0.75 | 0.76 | 0.74 | 0.74 | 0.71 |
| PF00636 | 0.73 | 0.85 | 0.84 | 0.81 | 0.78 | 0.71 |
| PF01368 | 0.73 | 0.84 | 0.83 | 0.79 | 0.77 | 0.72 |
| PF05226 | 0.79 | 0.8 | 0.81 | 0.8 | 0.8 | 0.78 |
| PF02811 | 0.83 | 0.95 | 0.92 | 0.88 | 0.86 | 0.81 |

Table S5: $ESS_r$ using salient and bottom neurons for PT-embed

| **Dataset** | **All** | **Top** | | | | **Bottom** |
|---|---|---|---|---|---|---|
| | | **(10%)** | **(25%)** | **(50%)** | **(75%)** | **(25%)** |
| PF00189 | 0.79 | 0.84 | 0.84 | 0.83 | 0.82 | 0.75 |
| PF00366 | 0.80 | 0.79 | 0.82 | 0.83 | 0.82 | 0.79 |
| PF02617 | 0.82 | 0.86 | 0.88 | 0.86 | 0.83 | 0.79 |
| PF00934 | 0.88 | 0.85 | 0.88 | 0.88 | 0.88 | 0.85 |
| PF06130 | 0.97 | 0.98 | 0.98 | 0.98 | 0.98 | 0.93 |
| PF12704 | 0.68 | 0.69 | 0.71 | 0.7 | 0.7 | 0.66 |
| PF13423 | 0.73 | 0.8 | 0.77 | 0.75 | 0.75 | 0.72 |
| PF05226 | 0.76 | 0.77 | 0.8 | 0.78 | 0.78 | 0.74 |
| PF01368 | 0.76 | 0.78 | 0.82 | 0.82 | 0.79 | 0.72 |
| PF02811 | 0.86 | 0.91 | 0.89 | 0.89 | 0.89 | 0.81 |

Table S6: $ESS_r$ using salient and bottom neurons for MSA-embed

| **Dataset** | **All** | **Top** | | | | **Bottom** |
|---|---|---|---|---|---|---|
| | | **(10%)** | **(25%)** | **(50%)** | **(75%)** | **(25%)** |
| PF20554 | 0.73 | 0.8 | 0.79 | 0.79 | 0.76 | 0.7 |
| PF02617 | 0.74 | 0.89 | 0.85 | 0.81 | 0.78 | 0.69 |
| PF00934 | 0.78 | 0.82 | 0.84 | 0.83 | 0.82 | 0.75 |
| PF03862 | 0.78 | 0.94 | 0.89 | 0.9 | 0.86 | 0.72 |
| PF06130 | 0.79 | 0.93 | 0.93 | 0.91 | 0.88 | 0.72 |
| PF00636 | 0.65 | 0.63 | 0.67 | 0.68 | 0.68 | 0.6 |
| PF04427 | 0.69 | 0.79 | 0.8 | 0.76 | 0.71 | 0.66 |
| PF01885 | 0.71 | 0.79 | 0.82 | 0.8 | 0.76 | 0.67 |
| PF04051 | 0.72 | 0.82 | 0.83 | 0.8 | 0.77 | 0.66 |
| PF00006 | 0.72 | 0.81 | 0.85 | 0.81 | 0.78 | 0.69 |

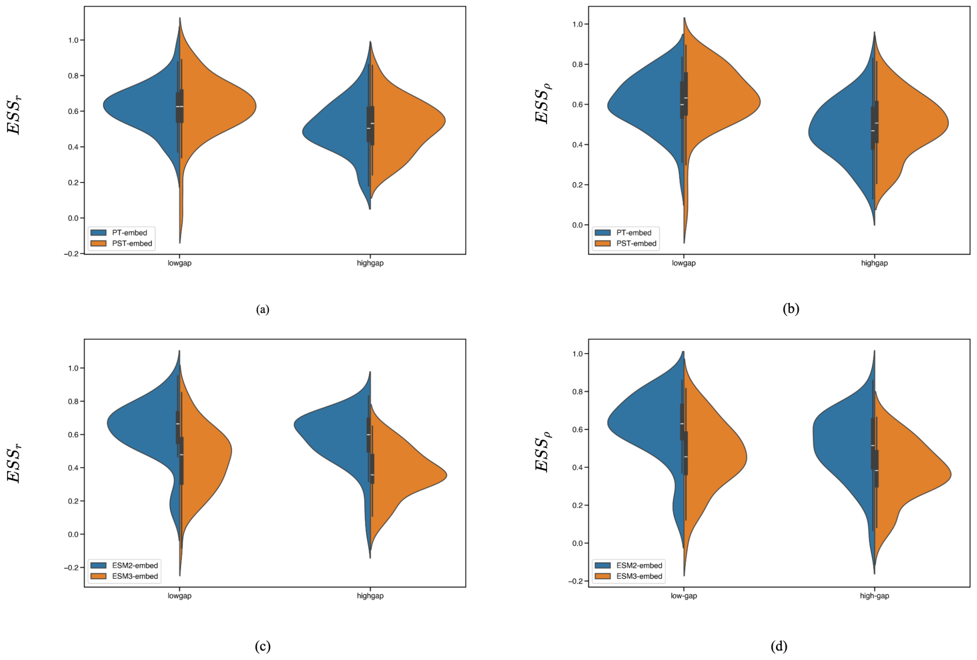


Fig. S8: Distributions of $ESS_r$ of PT-embed vs PST-embed (a) and ESM2-embed vs ESM3-embed (b) across all applicable low-gap and high-gap datasets. Distributions of $ESS_\rho$ of PT-embed vs PST-embed (c) and ESM2-embed vs ESM3-embed (d) across all applicable low-gap and high-gap datasets.

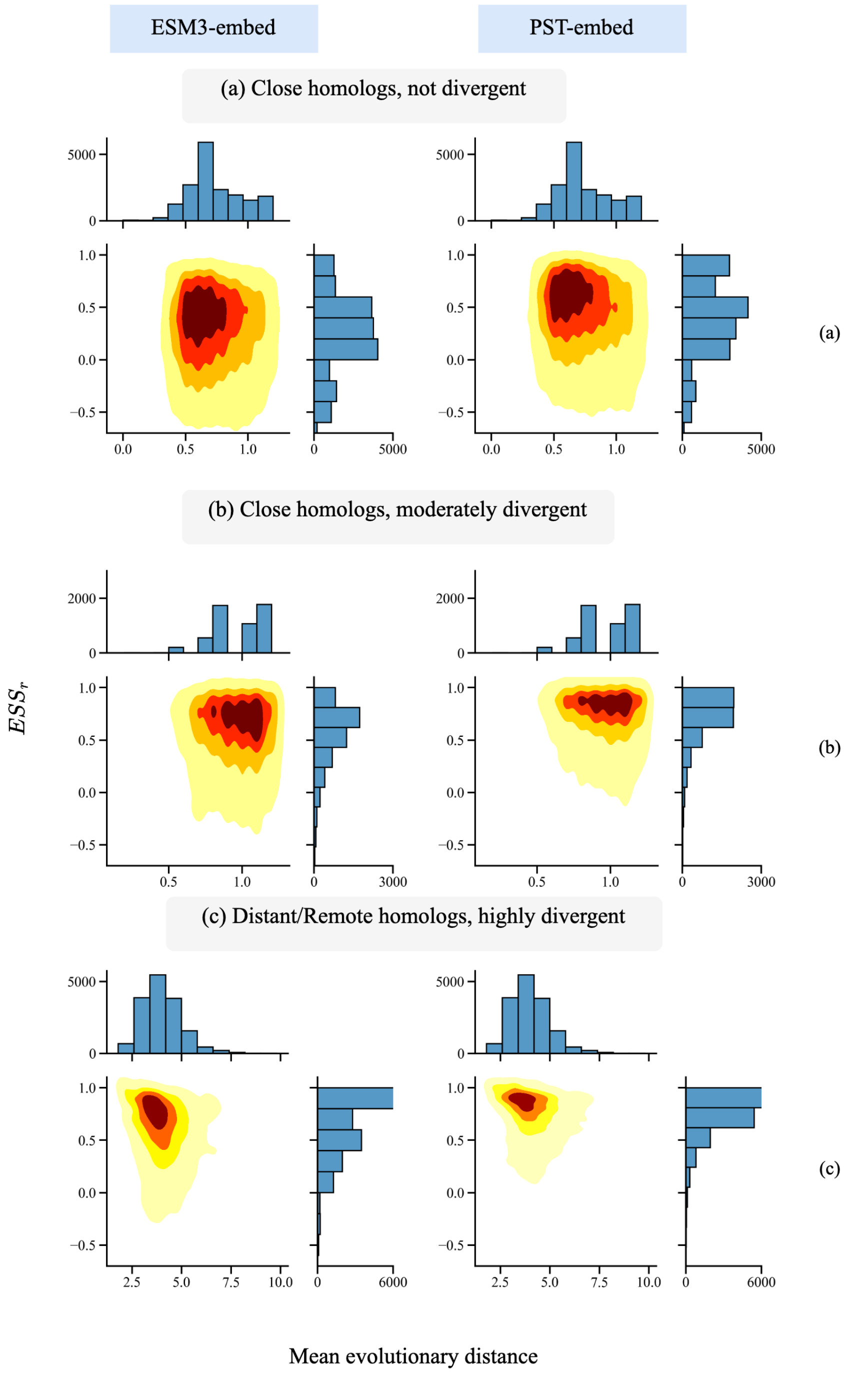


Fig. S9: Analysis of $ESS_r$ for different evolutionary distances for ESM3-embed and PST-embed. Density of $ESS_r$ for "close homologs, not divergent" (a), "close homologs, moderately divergent" (b) and "distant/remote homologs, highly divergent" (c) relative to mean evolutionary distance in each group (from reference sequence).

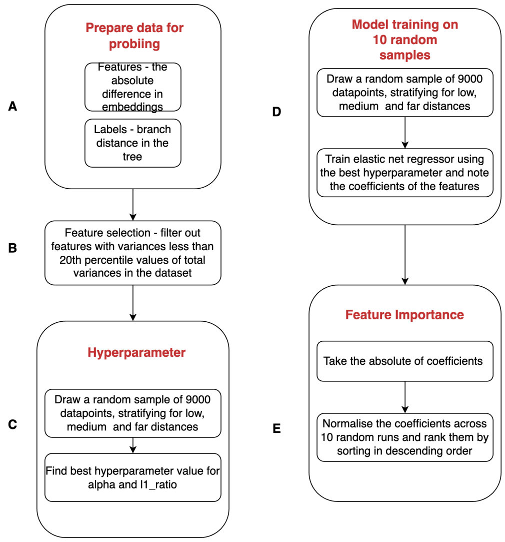


Fig. S10: Evolutionary probe workflow. (a) Data preparation. (b) Feature selection. (c) Hyperparameter selection. (d) Model training. (e) Feature importance.
